# 4sUDRB-sequencing for genome-wide transcription bursting quantification in breast cancer cells

**DOI:** 10.1101/2020.12.23.424175

**Authors:** William F. Beckman, Miguel Ángel Lermo Jiménez, Perry D. Moerland, Hans V. Westerhoff, Pernette J. Verschure

## Abstract

Epigenetics maintains cell-identity specific gene-expression patterns. However, within a population of isogenic cells of the same identity, a substantial variability in gene expression and responsiveness is still observed. Transcription bursting is a substantial source of this gene-expression variability or ‘noise’, contributing to phenotypic heterogeneity and potentially driving both physiological and pathological processes such as differentiation or tumorigenesis and drug resistance. Identification of transcription-bursting dynamics at a genome-wide scale has been restricted to inferring bursts in mRNA production computationally from the heterogeneity of mRNA levels in single cell transcriptomic data. Systematic characterisation of the genomic and epigenetic chromatin context of genes with defined transcription bursting behaviour has been incomplete. Here, we measured the bursting of transcription itself by genome-wide nascent RNA sequencing of breast cancer MCF-7 cells upon synchronisation of transcription with a transcription elongation inhibitor and by calibration using live cell imaging of nascent PP7-tagged *GREB1* transcription. Comparing across the entire genome, we find transcription bursting to be ubiquitous, with burst sizes of up to 160 transcripts. Transcription bursting is strongly correlated with steady state gene expression between genes, whereas both burst frequency and nascent transcript degradation only correlate weakly. Individual genes deviate strongly from this trend and engage both in anomalous burst size and frequency. We find that the presence of the TATA box or Inr sequence within gene promoters are significantly associated with a larger burst size, as are promoter-associated YY1 and E2F1 transcription-factor binding motifs. Enrichment of the transcription start site with epigenetic marks such as H3K79me2 and H3K27ac is also strongly associated with the transcription burst size. Finally, we show that in these MCF-7 breast-cancer cells, genes with a larger transcription burst size exhibit a larger immediate transcriptional response following endocrine drug treatment. Our genome-wide transcription-bursting analysis method paves the way to elucidate the dynamic role of epigenetic regulation on dynamic transcription in pathophysiology.

## Introduction

Gene expression noise is ubiquitous in both unicellular and multicellular life, permitting population survival in new environments and initiation of phenotypic divergence. Accordingly, gene expression noise has become a focus for multiple branches of biology, particularly those addressing emergence of complex traits such as drug resistance. A major part of this noise stems from the intrinsic propensity for RNA production to occur during short irregular bursts of promoter activity interspersed by longer periods of relative promoter inactivity ^1^, with the resulting noise depending on the number of RNA molecules produced during the promoter’s ‘ON’ state (burst size), the frequency of transitions to the ON state (burst frequency), and the degradation rate of the resulting transcript. This mode of RNA production, termed transcription bursting, is widespread in metazoans, protozoans, plants and bacteria ^2–6^.

In the absence of multiple promoter states, RNA production would still be somewhat noisy, resulting in a Poisson distribution of gene expression across any population of cells. In this case the variance in gene expression across the population should equal the mean expression level and the average behaviour of the cell population should reflect the average expression level. However, if a gene promoter is capable of switching between multiple states of activity, the resulting sporadic transcription bursts in mRNA production would lead to an expression distribution with increased variance compared to the mean level, referred to as ‘overdispersion’ ^7,8^. If propagated to the functional level, such overdispersion would then cause the cell population to behave differently on average with outliers determining functionality more than the average.

Transcription bursting and the associated noise in gene expression have been implicated in phenotypic bet hedging in both unicellular organisms ^9^ and in cancer ^10^, where they serve to increase the survival probability of a population of cells when facing dynamic environmental stress. In cancer, high transcriptional variability is associated with poor prognosis and resistance to therapy ^11–14^. Drug resistance emerges in 30-40% of patients during endocrine therapy of estrogen receptor (ERα)-positive breast cancer and is a main barrier to successful cancer treatment ^15^. Indeed, *in vitro* studies have suggested that breast cancer cells are capable of freely transitioning between several phenotypic states including a putative cancer stem-cell phenotype ^16^, suggesting that cell states are susceptible to non-genetic fluctuations. *In vitro* studies of melanoma have revealed that transcription bursting can give rise to a transient small population (∼1 in 3,000) of pre-resistant cells based on concurrent high expression of several genes involved in resistance pathways ^11,17^. Non-genetic heterogeneity has directly been shown to contribute to acquired drug resistance via inheritance of acquired characteristics in tracked single cells ^18^.

It is therefore important to determine which transcription features are responsible for the non-Poisson noise in mRNA and protein levels. With this in mind, pre-determined models of transcription bursting have been fitted to expression distribution data generated by single cell techniques such as single molecule RNA fluorescent *in situ* hybridisation (RNA FISH) and single cell transcriptomics, with the goal of inferring transcription bursting parameters ^19^. These methods of transcriptionally profiling single cells are static and, either extremely low throughput (RNA FISH) or subject to inaccuracy and technical biases for any transcript that is not highly abundant (single cell transcriptomics). Additionally, any estimated parameters depend on the pre-assumed model and often the expression distribution can computationally be fitted with multiple different models of transcription bursting or with models of static genetic heterogeneity within a cell line. Conversely, transcription bursting dynamics can be quantified using live cell reporters of transcription, but this relies on the integration of MS2- or PP7-binding loops at the locus of interest ^20,21^. This gene specific approach does not enable genome-wide quantification of transcription bursting. Currently no experimental method exists for characterizing transcription bursting dynamics of endogenous genes at a genome-wide scale.

In order to facilitate the study of transcription bursting and the molecular mechanisms that govern it, we present a genome-wide burst-size quantification technique based on 4-thiouridine (4sU) sequencing upon discontinuation of the cells’ exposure to the transcription elongation inhibitor 5,6-Dichloro-1-β- d-ribofuranosylbenzimidazole (DRB). By synchronising the MCF-7 breast cancer cells through quick removal of DRB and concomitant incubation with 4sU ^22,23^, we are able to use bulk RNA sequencing of 4sU-labelled nascent transcripts to quantify the magnitude of the nascent transcriptional response. While these kinds of global run-on experiments have classically been used to measure both RNA polymerase elongation and RNA degradation rates, or to identify novel transcription start sites, we instead use this methodology to quantify the magnitude of the nascent transcriptional response. Using a PP7 live cell reporter system we show that DRB-induced synchronisation of transcription does not affect the burst size measurements and that the peak in transcription following DRB removal accurately quantifies the transcription burst size. Our study points out that genes with larger transcription burst sizes are significantly enriched for YY1 and E2F1 transcription factor binding motifs at their promoter and we find multiple associations of transcription bursting with specific histone modifications and core promoter elements. Our findings are not only essential to fundamentally understand genome-wide transcription dynamics but may also constitute a prelude to facilitating the modulation and/or prediction of transcriptional noise in a clinical setting.

## Results

### Nascent *GREB1* transcription bursts in individual cells

The mRNA content of single cells fluctuates over time with these fluctuations being proportional to the transcription burst size ^24^. Bulk cell measurements of transcription obscure these fluctuations because there is a lack of synchronicity between individual cells. When cells are synchronised, for instance through transcriptional inhibition and subsequent release thereof, a short-lived wave of RNA production can result which is followed by a synchronous transition of promoters from an ON state to an OFF state and a gradual decay of the RNA ^25,26^. This is because gene promoters recently subject to an ON-to-OFF state transition are refractory towards re-stimulation ^27^. In support of this, we have previously shown that, in the absence of oscillations in concentrations of transcription factors around the promotor, the initial peak of transcription can only robustly occur given multiple promoter activity states corresponding to a clock mechanism of transcription activation ^28^. The amplitude of this peak should be directly proportional to the transcription burst size of a gene during steady state gene expression and should reflect the single cell dynamics of transcription.

To examine transcription dynamics following transcriptional synchronisation, we used an MCF7□GREB1□PP7 cell line containing 24 PP7 loops integrated into the second exon of the endogenous *GREB1* locus (Supplementary Figure 1) ^29^. Stable expression of the GFP-labelled PP7-binding coat protein (PCP-GFP) allowed for direct visualisation of the accumulation of nascent transcripts as bright GFP nuclear foci, presumably indicating the sites of transcription (Figure 1A). In order to synchronise transcription between the cells, we treated the cells with 5,6-Dichloro-1-β-d-ribofuranosylbenzimidazole (DRB) a potent yet reversible inhibitor of the CDK9 kinase subunit of P-TEFb. DRB thereby inhibits RNA elongation by preventing serine 2 phosphorylation on the C-terminal domain of RNA polymerase II ^30^. By adding GFP at a known concentration to the imaging media we quantified the transcripts at every transcription site. After 3 hours of DRB treatment we measured on average 0.9 transcripts at each transcription site compared to 4.1 transcripts in the cells treated with DMSO as vehicle control (Figure 1B, C). These measurements suggest an inhibitor efficacy of over 75%. Confocal imaging of individual living cells revealed bursts in transcription across individual cells (Fig. 1B, Supplementary Figure 2), independently of the pre-incubation with DRB. After the pre-incubation with DRB, a more synchronous burst after DRB release is noted (Supplementary Figure 2) as compared with the pre-incubation with the vehicle control (DMSO), suggesting that our aim of using DRB to synchronize the cells’ transcription had been successful. Synchronicity should enable us to read the burst size from the maximum population average, which we found to be ∼10 transcripts per transcription site at approximately 50 minutes after transcription initiation (Figure 1C), which was followed by a refractory period of an hour. The burst size is calculated by taking the height of the first peak in transcription after DRB release minus the starting level, i.e. approximately 11.3 – 1 = ∼10.3. Such a strong pulse of ensemble-average transcription was not observed in the control cells that had been mock treated with DMSO solution only followed by substitution with DMSO free medium (Figure 1B-C). Instead, in these control cells a much weaker and earlier transient increase was found to occur, possibly as a result of providing the cells with fresh culture medium, or due to the binding of DMSO to dsDNA, which is known to interfere with polymerase activities ^31^.

**Figure 1.**
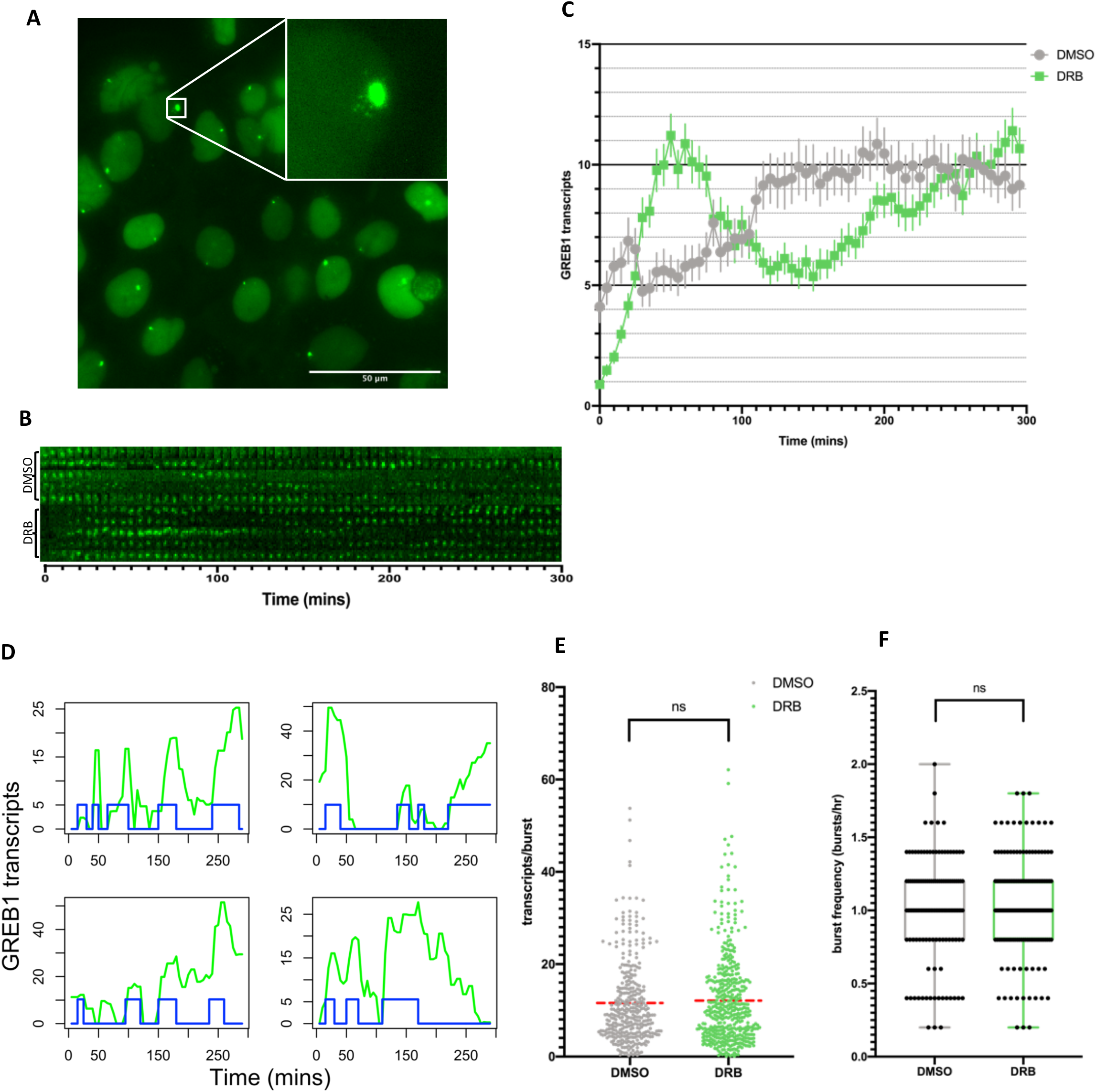
Single cell imaging confirms bursts of transcription in individual living cells that can be synchronised by DRB removal. **(A)** Representative image of PP7-tagged *GREB1* transcripts in MCF-7 GREB1-PP7 reporter cell line imaged with confocal microscopy. Transcription sites occur as bright GFP positive foci in the cell nuclei. Insert shows one transcription site, with several individual mRNAs that appear to translocate away from the site. **(B)** Representative time lapse imaging of PP7-tagged *GREB1* transcription sites showing images of PP7-GFP tagged transcription sites throughout the time course. The top five rows show transcription sites in DMSO-treated control cells and the bottom five rows show sites in DMSO+DRB transcription-synchronised cells. Images were taken at 5 minute time intervals. **(C)** Mean number of PP7-tagged *GREB1* transcripts at transcription sites following DRB synchronisation and release. n=132 for DMSO-only control (grey) and n=161 for DRB (green) from 3 biological repeats. Error bar = 2 times SEM. **(D)** Detection of transcription bursts events (blue) illustrated for 4 individual PP7 tagged transcription sites in DMSO control cells. Transcription site trajectories (green), show various burst sizes and frequencies. **(E)** Transcription burst sizes for DMSO treated (grey) and DRB treated (green) cells, calculated as the difference in transcript number between the beginning of each ON period (as in D) and the maximum number of transcripts detected for that same period. There was no significant difference between the two groups (Mann Whitney test p-value = 0.65). Red dotted lines indicate mean transcription burst size for each group. **(F)** Transcription burst frequencies for DMSO treated (grey) and DRB treated (green) cells, calculated as the number of detected peaks per hour of the time course. No significant difference was found between the two groups (Mann Whitney test p-value = 0.84).

To assess the levels of transcriptional heterogeneity in response to DRB treatment across single cells, we detected the transcription peaks in the individual cells during their time lapse measurements (Figure 1D, Supplementary Figure 3). This allowed for the quantification of transcription burst sizes and frequencies of individual cells during the 5 hour imaging period. The heterogeneity was extremely high, in terms of both burst frequency and burst size (Figure 1E, F), with some cells producing 60 transcripts in a single burst, but others fewer than 5 transcripts. Based on these individual cell transcription bursting measurements, the average burst size across the 5 hours in which the cells were analysed was 12 transcripts, in good accordance with the cell-population average of 10 transcripts determined within the first wave of average transcription (Figure 1C). Comparison of the time-lapse transcription measurements upon DRB synchronisation to the DMSO vehicle control traces (Figure 1E, F) shows that neither the transcription burst sizes nor the burst frequencies of the individual cells were significantly affected by the DRB treatment, confirming that DRB synchronisation is suitable for our transcription measurements in synchronised cells.

### 4sUDRB-seq to measure genome-wide transcription bursting

Whilst these results were in terms of actual mRNA numbers, they were limited to a single gene, i.e. PP7-tagged *GREB1*. We next assessed transcription burst sizes genome-wide using 4sUDRB sequencing (Figure 2A). In 4sUDRB-seq, cells are allowed to transcribe in the presence of 4-Thiouridine (4sU) for up to 40 minutes after the removal of DRB. The time-dependent monitoring of 4sU incorporation into nascently transcribed RNA provides a view of the underlying transcriptional dynamics at the minutes’ timescale. Upon 3 hours of DRB treatment transcription elongation was again stalled with subsequently fewer mapped reads than for DMSO control treated cells, particularly in exonic regions, while DRB removal allowed RNA elongation by RNA Polymerase II (RNAPII) (Figure 2B). The promoter-proximal number of mapped reads declined starting at 10 minutes after DRB release, whereas production of RNA further along the gene still increased; suggesting a transient increase (‘peak’) of promoter-proximal transcriptional activity.

**Figure 2.**
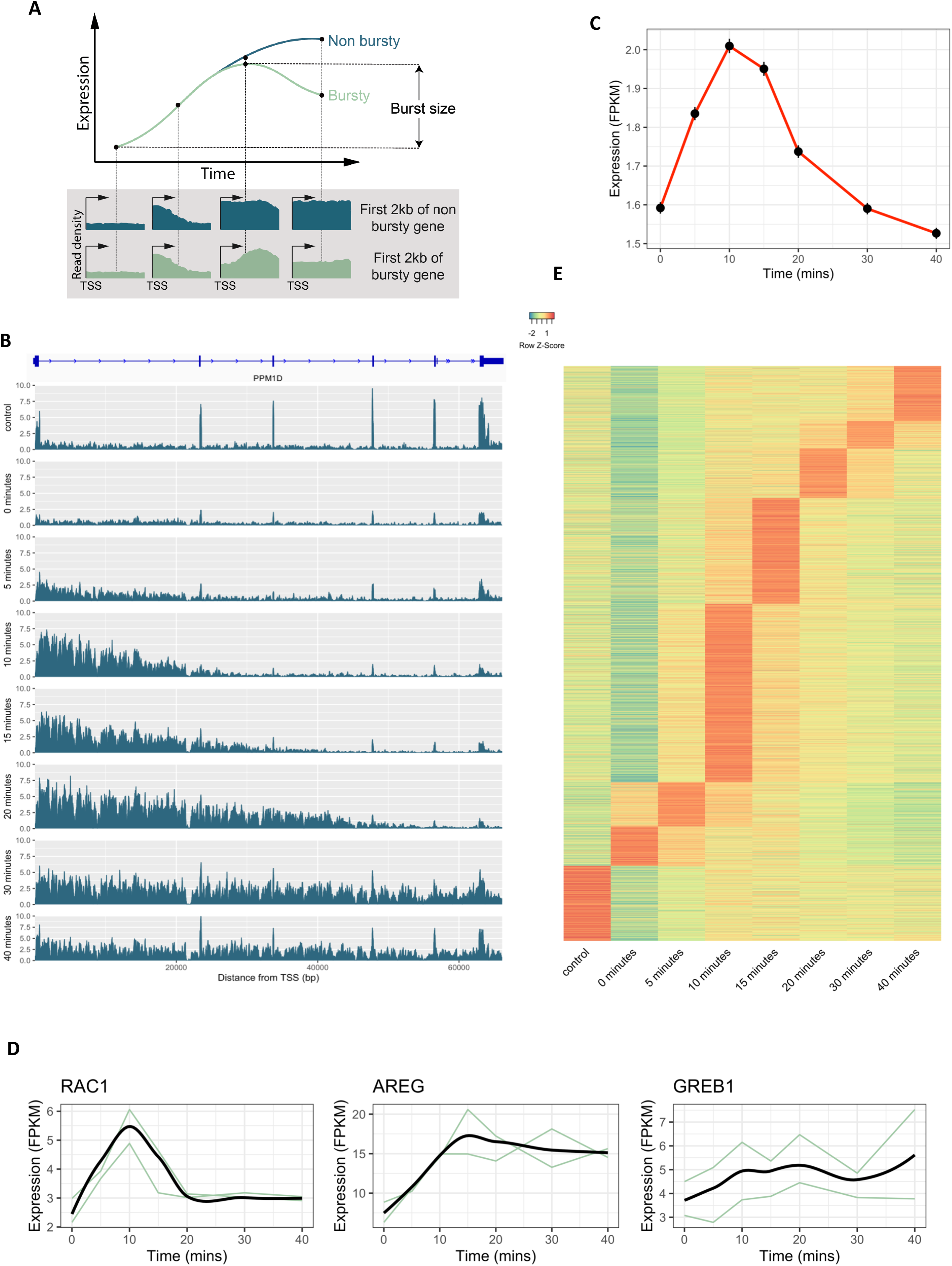
4sUDRB sequencing reveals that the majority of genes are transcribed in bursts. **(A)** Scheme illustrating burst size quantification from 4sUDRB sequencing data. RNA expression for genes not transcribed in bursts (blue) will increase monotonically to their steady state gene expression plateau. Any gene transcribed in bursts (green) will show a peak in RNA production followed by a decay in later time points. The burst size is here quantified as the height of this initial peak. Bottom panel shows the read densities expected at four time points after removal of the transcription inhibitor as a function of distance from TSS for the regions directly following the transcription start sites of non-bursty (blue) and bursty (green) genes. **(B)** An example of normalised 4sUDRB-seq read depth versus the distance from the TSS in base pairs along the *PPM1D* gene Panels are ordered by increasing time, with the DMSO control sample at the top followed by the 4SUDRB samples. The y-axis shows the read depth normalised by total read counts. **(C)** Mean gene expression values for the 11,466 genes that passed low expression and low length filtering; error bar = 2 times SEM. FPKM values were calculated using the first 2 kb region following transcription start sites, with exonic regions subtracted. **(D)** 4sUDRB-seq time-course results for three genes (*RAC1*, *AREG* and *GREB1*) with both biological replicates (green) and loess smoothed means (black). Each of the genes use a different y-axis **(E)** Heatmap of 4sUDRB-seq expression time courses (in FPKM) for the first 2kb of 11,466 genes during the 40 minutes of transcription. The time-course trajectory for each gene was smoothed by fitting a local polynomial regression (locally weighted scatterplot smoothing, LOESS) and genes were grouped based on the time point at which maximum expression occurred. The control sample was not included in the LOESS smoothing.

To examine whether this nascent transcription peak was a genome-wide phenomenon, we quantified the number of RNAs (‘read depth’) within the first 2 kb of intronic regions of every gene throughout 40 minutes of 4sU incorporation. For the 11,466 genes that passed low-expression and small-size filtering (see Methods), we observed a peak in average number of nascent RNAs at 10 minutes after DRB release. This elevated expression disappeared again by 40 minutes (Figure 2C). This peak was small because individual genes varied significantly in their nascent average transcriptional response. For example *RAC1* showed a rapid initial spike in transcription followed by a rapid decay in RNA, whereas the decay of *AREG* was much slower (Figure 2D). *GREB1* also showed a peak in transcriptional activity, with the largest number of mapped reads at 20 minutes after DRB removal which subsequently resides presumably as the intronic RNA is rapidly co-transcriptionally spliced and degraded (Figure 2D). This peak in transcription occurs earlier than the peak detected in PP7 based live cell imaging because we focus on the first 2 kb of each gene, while PP7 integration was ∼15 kb further into the gene. Grouping genes based on the time at which their expression peak occurs revealed that the transcription peak occurred most frequently at 10 minutes (31% of genes) followed by 15 minutes (18% of genes) (Figure 2E). The vast majority of genes (80%) show a maximal expression that exceeds the expression in the control and 0 minutes samples, revealing that these genes produce more RNA in the nascent transcriptional wave than at steady state. Only a small subset of genes (13%) did not regain steady state levels of gene expression during the 40 minutes following DRB release. As expected, using a window size of less than 2kb to quantify expression resulted in an earlier peak of transcription, and using a larger window size resulted in the peak occurring later. Varying the window size resulted in a plateau for mean burst sizes around the 2kb window size (Supplementary Figure 4).

### 4sUDRB-seq calibration to determine transcription burst sizes genome-wide

Genes that failed to respond upon DRB release (i.e. having maximal expression at t=0 or showing no increase in expression between t=0 and t=5) were omitted from further analysis (Figure 3A). Additionally, any genes which did not show a decay of at least 10 % of read depth at any point in the 40 minutes following DRB release were ignored since they appeared to remain in the ON state. For the remaining 8,429 genes burst sizes were quantified. The burst sizes for these remaining genes were calculated based on the difference between read counts at 0 minutes and the maximal read count following DRB release (Figure 2A). In order to convert the burst size from FPKM to absolute transcript numbers, the *GREB1* burst size of 10.33 transcripts based on PP7 reporter gene measurements was used as a calibration factor. The *GREB1* 4sUDRB-seq burst size is 1.8 FPKM, and so the conversion factor between FPKM and transcripts is equal to 5.7.

**Figure 3.**
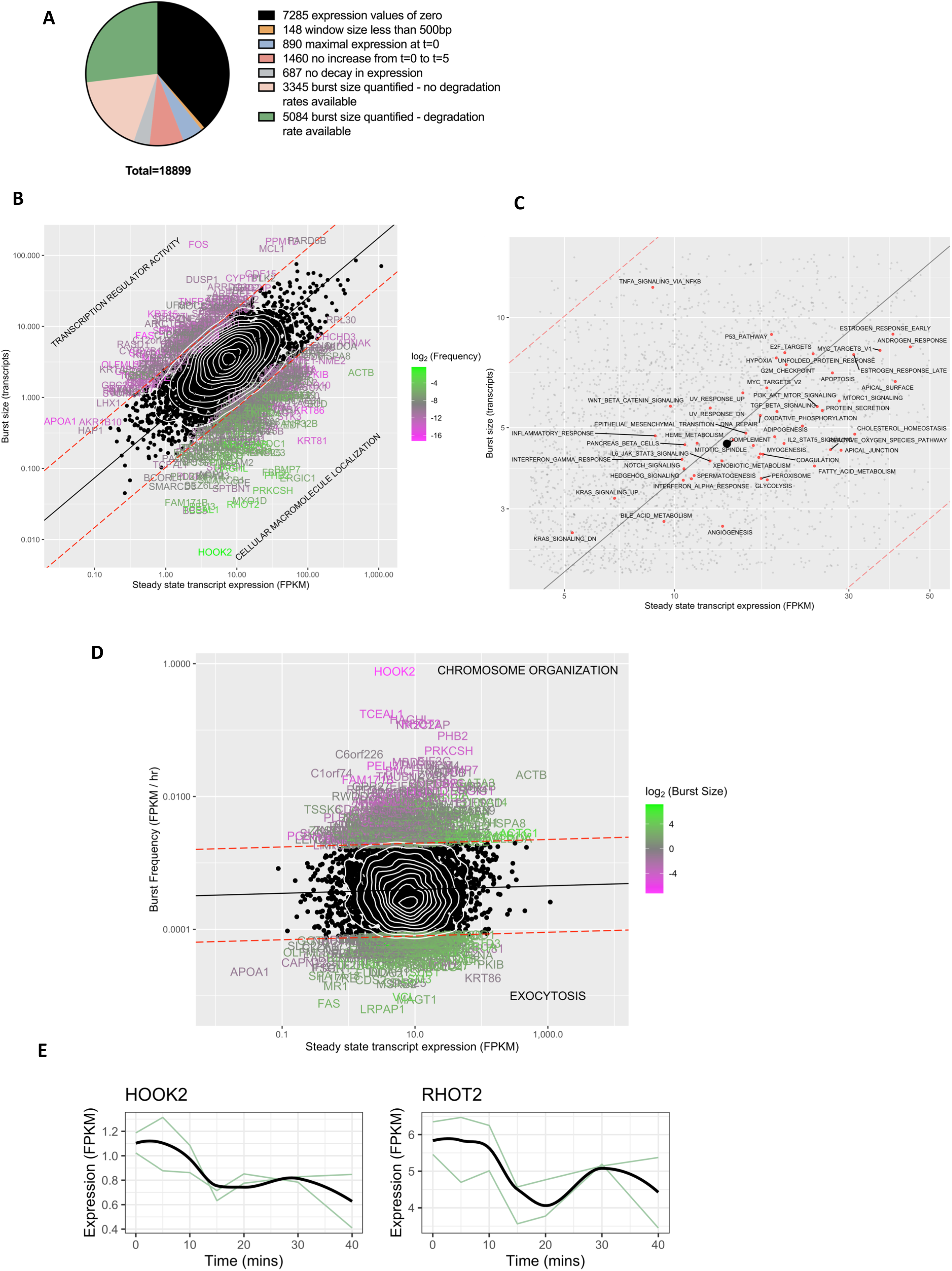
Transcription burst size and not burst frequency correlates with the steady state gene expression level. **(A)** Pie chart summarising categorisation of the 18,899 protein coding genes analysed. Figure 2 uses 11,466 genes that passed low expression and low size filtering (blue, red, grey, pink and green), while subsequent analysis used the 5,084 genes in the green group unless stated otherwise. **(B)** Deming regression (black solid line) of steady state gene expression level and transcription burst size for 5,084 genes. Genes are coloured by the calculated transcription burst frequency if their burst size exceeds a 5 fold deviation in either direction (red dotted lines) from the value predicted by the regression. The top overlapping GO term for each group is also shown. **(C)** Plot shows a zoomed in section of (B) with individual genes (small black dots) and the average of all 5,084 genes (large black dot). The mean of each MSigDB hallmark gene set is plotted (red dots). Diagonal lines represent the same trend lines as (B). TNFA signalling via NFKB genes on average has a much larger transcription burst size than expected, while genes involved in various metabolic pathways have much smaller burst sizes than expected for their expression level. **(D)** Deming regression (black solid line) of steady state gene expression level and transcription burst frequency for 5,084 genes. Genes are coloured by burst size if their calculated transcription burst frequency exceeds a 5 fold deviation in either direction (red dotted lines) from the value predicted by the regression. The top overlapping GO term for each group is also shown. **(E)** 4sUDRB-seq time course results for two genes with the highest transcription burst frequency (*HOOK2* and *RHOT2*) with both biological replicates (green) and LOESS smoothed means (black). The two genes use different y-axes.

Genes with the largest burst sizes can be found in Table 1. The largest burst size of 162 transcripts was calculated for *PARD6B* (Figure 3B). The *TFF1* burst size was estimated to be 71 transcripts per burst, in good accordance with previously observed measurements based on *TFF1* RNA FISH ^32^. For the whole gene set, the average burst size was 3.7 transcripts, but individual burst sizes were highly variable, with a standard deviation between genes of 5 transcripts.

**Table 1.** Transcription parameters for the genes with the largest transcription burst size from 4sUDRB-seq experiments. Transcript degradation rates were taken from ^33^. Steady state gene expression (ssExp) was calculated from the control sample of the 4sUDRB-seq data.

| Genes | Burst size<br>(transcripts) | kdeg<br>(1/hour) | Burst frequency<br>(FPKM/hour) | ssExp<br>(FPKM) |
| --- | --- | --- | --- | --- |
| PARD6B | 162 | 0.010757 | 0.001500 | 97.5 |
| PPM1D | 159 | 0.002800 | 0.000164 | 43.6 |
| FOS | 143 | 0.008835 | 0.000066 | 2.9 |
| MCL1 | 125 | 0.004581 | 0.000447 | 31.3 |
| HSPB1 | 85 | 0.000546 | 0.001719 | 475.7 |
| APPBP2 | 75 | 0.001631 | 0.000256 | 69.2 |
| TFF1 | 71 | 0.000263 | 0.000257 | 1086.0 |
| AREG | 60 | 0.001429 | 0.000348 | 71.4 |
| MYL12A | 57 | 0.000583 | 0.000086 | 120.5 |
| DDX5 | 55 | 0.001515 | 0.000811 | 101.5 |
| GDF15 | 55 | 0.002759 | 0.000167 | 21.7 |
| PSMB4 | 52 | 0.000997 | 0.001333 | 151.3 |
| PLK2 | 48 | 0.010981 | 0.002389 | 24.1 |
| CYP1B1 | 48 | 0.000793 | 0.000137 | 12.7 |
| DUSP1 | 46 | 0.022255 | 0.000734 | 3.2 |
| RPS6KB1 | 45 | 0.001457 | 0.000280 | 70.1 |
| TRIM37 | 45 | 0.001401 | 0.000334 | 190.9 |
| CSDE1 | 45 | 0.000728 | 0.000195 | 264.7 |
| ACTG1 | 45 | 0.000554 | 0.002825 | 366.3 |

### 4sUDRB-seq calibration to estimate transcription burst frequencies genome-wide

Our genome-wide 4sUDRB-seq calibration method does not enable direct measurement of transcription burst frequencies (unless the frequency is very high). Yet, from the non-DRB treated control cells the steady state gene expression level can be determined, and from the DRB treated cells the burst size can be determined, both genome wide. In a deterministic situation (i.e. when every gene has a defined burst size and frequency), the expression level of a gene should equal the arithmetic product of transcription burst size and burst frequency divided by the mRNA degradation rate constant. Consequently: *f* = *k_d_* · *f*′ = *k_d_* · *nRNA_ss_*/*b*, where *f, b, k_d_* and *nRNA_ss_* refer to the transcription burst frequency, burst size, nascent mRNA degradation rate, and the nascent mRNA expression at steady state. With *f* we refer to the transcription burst frequency normalised by the degradation rate constants taken from Schueler et al.^33^.

### Expression-correlation of transcription burst size but not burst frequency is a genome-wide trend

In order to examine whether, across the genome, gene expression was determined by transcription burst size, by burst frequency, or by both, we analysed the 5,084 genes for which transcript-degradation rates were available (see above for the reason of this restriction). We found a complex answer to this question: on the one hand, the transcription burst size correlated positively with the mean mRNA expression level, with a slope in the log-log plot not too distant from 1 (≈0.84; Figure 3B). On the other hand, many individual genes deviated significantly (up to a factor of 5) from this trend line. Genes with transcription burst sizes larger than expected on the basis of the trend line and their mean expression level, were significantly associated with transcriptional regulator activity, whereas the group of genes showing a smaller transcription burst size than expected on the basis of the trend line were significantly associated with cellular macromolecule localisation. Genes located in the upper left corner are likely to be the most heterogeneous in gene expression, while those in the bottom right are expected to be the most homogeneous. Multiplying steady state expression levels by transcript degradation rates had minimal effect on the gradient of trend line or the residual error of genes from the trend line (0.84 to 0.85 and 0.33 to 0.32 respectively, Supplementary Figure 5). We plotted the centres of mass for each MSigDB hallmark gene set onto these axes to see if any groups of genes deviated from the trend line relationship between the transcription burst size and steady state gene expression level (Figure 3C). TNFA signalling via NF-kB was strongly shifted towards the upper left, suggesting that genes within this group are extremely heterogeneous in gene expression, with transcription burst sizes much larger than expected for the mean expression level. Other groups of genes that shifted similarly included WNT signalling and P53 signalling. Many hallmark gene sets involved in metabolism were shifted towards the bottom right (including glycolysis, fatty acid metabolism and cholesterol homeostasis), suggesting more homogeneous gene expression across the population.

We found neither positive nor negative correlation between the transcription burst frequency and the steady state gene expression level (Figure 3D), but again strong differences between individual genes are noted. The genes with high transcription frequency were most significantly associated with chromosome organisation, whereas genes with low frequency were most significantly associated with exocytosis. To validate the transcription burst frequency calculations, we evaluated the time courses of genes that exhibit high transcription frequency (*HOOK2* and *RHOT2*) revealing that they indeed undergo multiple transcription bursts already within the 40 minute time course (Figure 3E). As could be inferred from the above results, the RNA degradation rate constants correlated somewhat negatively with expression level (Supplementary Figure 6)

### Exon FISH monitoring of nascent mRNA at transcription sites

To confirm that most transcription was indeed occurring in transcription bursts, we performed RNA FISH for 8 genes that according to the 4sUDRB-seq results had transcription burst sizes mostly in excess of 3 transcripts, with various gene expression levels. RNA FISH analysis allows for visualisation of single RNA transcripts that occur dispersed throughout the cell, but also highlights accumulation of nascent transcripts in the nucleus as bright foci (Figure 4A). In this experiment we used probes hybridizing to exon sequences. To confirm that the larger bright foci are indeed transcription sites, we performed intron RNA FISH additionally (Figure 4B). As splicing occurs co-transcriptionally ^34^, the intron segments of the RNAs should be degraded at the transcription site and not venture to other positions in the cell. Indeed we found that virtually all intron signals co-localised with exon signals, but not *vice versa* (Figure 4B). By comparing the size and intensity of all exon RNA foci in the nucleus (Figure 4A), to the size and intensity of single mRNA molecules in the nucleus, we quantified the number of transcripts at each transcription site. Averaging the number of transcripts at transcription sites for each gene should not be expected to equal the transcription burst sizes precisely for several reasons. Firstly, for many of the sites, transcription may have only recently begun and so the number of RNA polymerases currently elongating may be a fraction of the true number. Alternatively, transcription may have recently terminated, allowing transcripts to diffuse away. We hypothesised that this effect would be more severe for shorter genes, where transcripts reside for a shorter time, while for longer genes multiple bursts may co-occur leading to an overestimation of the true transcription burst size. Secondly, single mRNAs that already diffused in the nucleus will be full length transcripts, while those currently undergoing synthesis at transcription sites will only be fractions of the full length. Neither of these issues apply to the 4sUDRB-seq calculated transcription burst sizes. To address these issues we normalised the data by taking the average number of transcripts detected at transcription sites for each gene, and adjusting the burst size by the ratio between the gene length to average gene length for the 8 genes analysed (see Methods).

**Figure 4.**
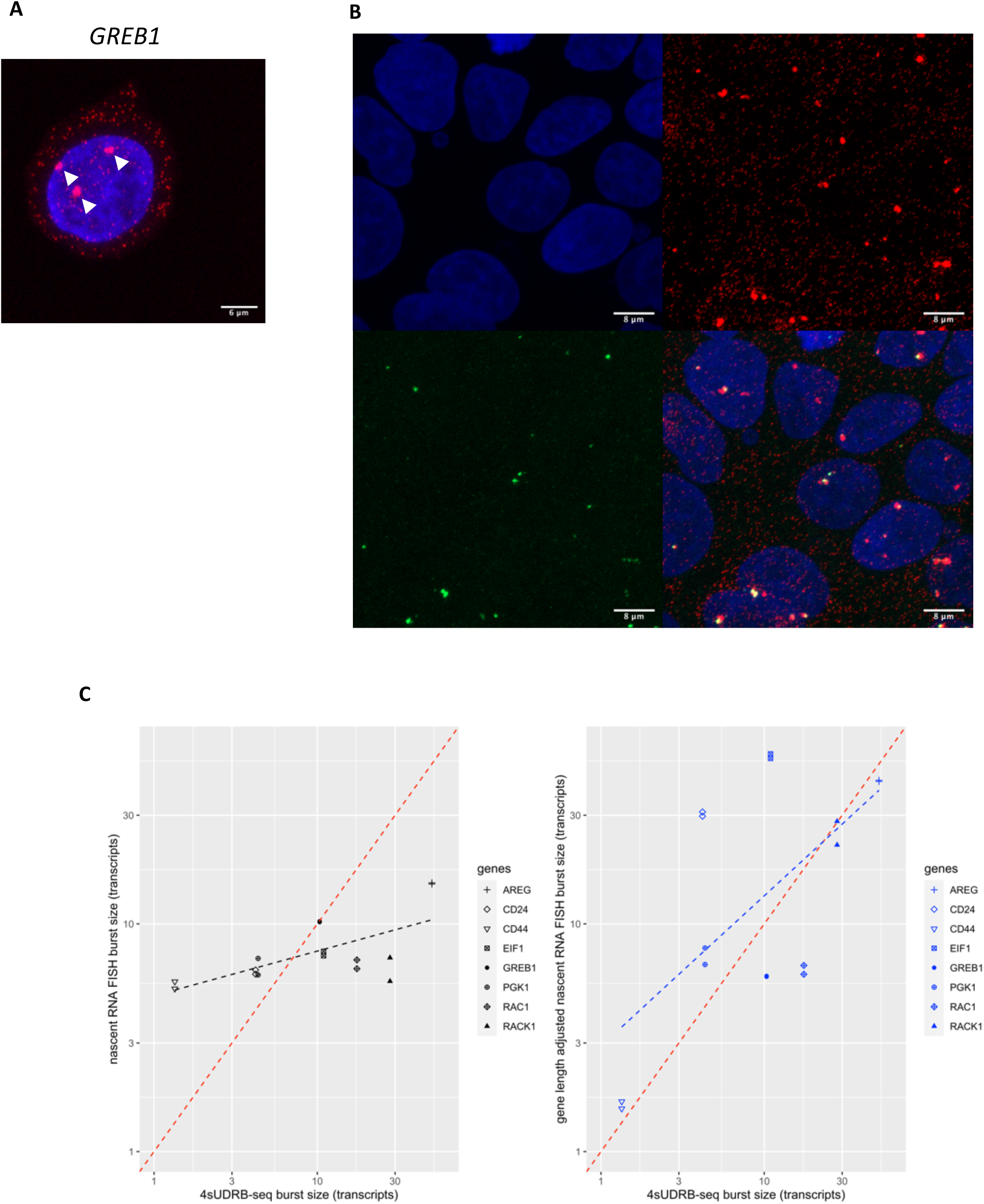
The extent of correlation between single molecule RNA fluorescence-*in-situ-* hybridisation burst sizes and 4sUDRB-seq burst sizes. **(A)** Representative image of an MCF-7 cell with *GREB1* transcripts labelled (red) following exon RNA FISH. Nuclei were counterstained with DAPI (blue). The small red spots show hybridisation with single RNA molecules, whilst the large transcription sites (at the white arrows) correspond to RNAs accumulated at transcription sites. **(B)** Representative image of MCF-7 cells with *GREB1* transcripts labelled following exon RNA FISH (red; upper right) and intron RNA FISH (green; bottom left) revealing frequent (but not universal) co-localisation of intron on exon signals (white spots in overlay: bottom right). Nuclei were counterstained with DAPI (blue; upper left). **(C)** Correlation between nascent RNA FISH-measured burst sizes of a set of genes (AREG, CD24, CD44, ElF1, GREB1, PGK1, RAC1, RACK1) with 4sUDRB-seq measured burst sizes. 4sUDRB-seq burst sizes were calibrated using the *GREB1* PP7 based burst size of 10.33 transcripts. Dashed red lines show what a perfect correlation should have led to (x=y). Left panel shows the raw nascent RNA FISH burst sizes plotted against the 4sUDRB-seq measured burst sizes. Each gene has two repeats, and each repeat has 100 cells. Some repeats overlap (*GREB1*, *AREG*). Black dotted line shows the linear regression trend line. Right panel shows nascent RNA FISH burst sizes of the respective genes adjusted based on gene length (see Methods). Dashed blue line shows linear regression trend line.

All of the 8 genes analysed had a transcription burst size higher than 1 transcript and 7 genes had a transcription burst size higher than 3 transcripts as determined by the 4sUDRB-seq experiment. The gene length-adjusted burst sizes from nascent RNA FISH also had burst sizes in excess of 1 transcript confirming that the 8 genes studied indeed produce transcription bursts. CD44 had the smallest burst sizes for both methods (1.3 transcripts versus 1.6 transcripts for 4sUDRB-seq and length adjusted RNA FISH respectively). The trend line of the gene length corrected burst size of the 8 genes was, perhaps remarkably, similar to the expected perfect correlation line (x=y) reflecting the 4sUDRB-seq determined burst sizes (Figure 4C, right panel). The individual genes however varied greatly, possibly due to the above considerations.

### *Cis*-acting regulatory features associated with transcription burst kinetics

The binding of transcription factors to specific *cis*-acting regulatory sequences within gene promoters has been suggested to be involved in regulating transcriptional burst kinetics ^35^. Therefore, we next wanted to test whether any transcription factor binding motifs are associated with particular transcriptional burst sizes or frequencies. We performed pre-ranked functional enrichment analyses using the GAGE ^36^ and FGSEA ^37^ methods. The only transcription factor significantly associated with a large transcription burst size by both methods was YY1 (Figure 5A), a transcription regulatory protein that is able to link enhancers with cognate promoters, and binds both DNA and RNA ^38,39^. The transcription factors of the E2F class were significantly associated with burst size based on the GAGE method, and appeared frequently in the FGSEA method but by itself this appeared not to be statistically significant. Hallmark gene sets significantly associated with a large transcription burst size according to both GAGE and FGSEA methods included estrogen response, G2M checkpoint and TNF signalling via NF-kB. (Supplementary Figure 7). To support the association of YY1 with an increased transcription burst size, we distinguished between genes with and genes without a YY1 binding motif (Figure 5B, C) anywhere between 1000 base pairs upstream and 100 base pairs downstream of the transcription start site (TSS). As expected, the average transcription burst size of genes containing this motif within that region was significantly larger than the average of all genes that did not contain this motif (p-value = 0.02, Figure 5C). Again, this reflects a trend across the genome; individual genes deviate from this average trend more often than not (Figure 5C). There was no significant difference in the mean expression level or the transcription burst frequency when comparing genes with and without the YY1 motif (Figure 5C). Similarly, when genes were discriminated based on the presence of a promoter motif for the transcription factor E2F1, genes containing this motif had a significantly larger burst size (p-value = 0.04), while there was no difference for mean expression or burst frequency (Figure 5C).

**Figure 5.**
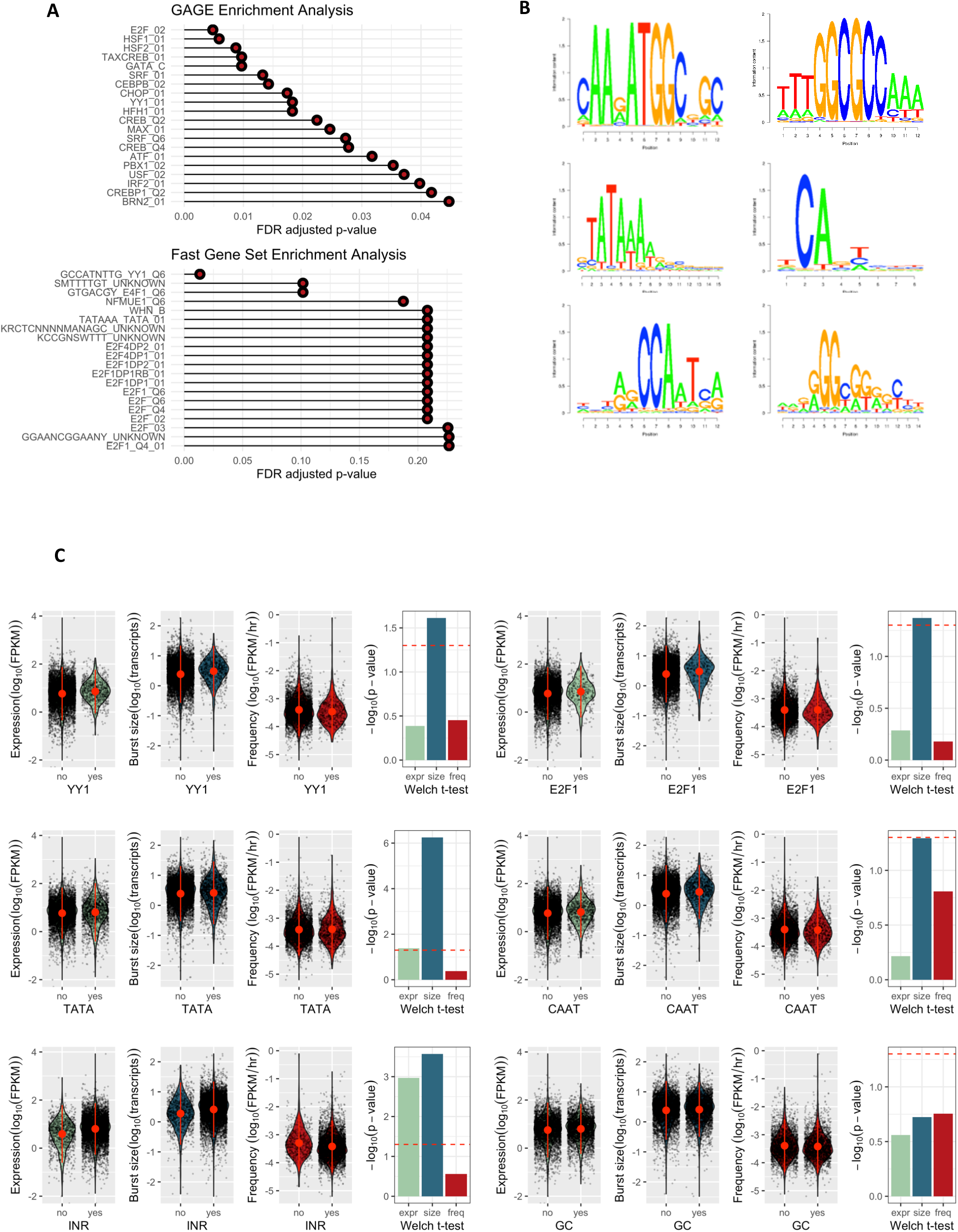
Increased transcription burst sizes are associated with promoter binding affinity of YY1 and E2F1 transcription factors and presence of promoter TATA box and Initiator elements. **(A)** Pre-ranked functional enrichment analysis for 5,084 genes using the GAGE (top) and FGSEA (bottom) methods. Pre-ranked gene lists were compared to the MSigDB transcription factor collection. Each analysis uses a different x-axis. **(B)** Motifs used (as taken from FindM) used to discriminate the genes. The order is the same as (C). **(C)** Violin plots of steady state gene expression (green), transcription burst size (blue) and transcription burst frequency (red) for genes grouped depending on presence (‘yes’) or absence (‘no’) of a promoter-proximal YY1 binding motif, E2F1 binding motif, TATA box, INR element, CAAT box and GC box. The significance between the two sets of genes in each violin plot is shown by the bar chart (Welch t-test). Red dotted line shows the p-value of 0.05.

Multiple studies have implicated a relationship between transcription burst size and specific core promoter elements ^19,40^, such as the TATA box, which recruits the TATA-binding protein (TBP) subunit of transcription factor II D (TFIID) and ultimately directs the formation of the transcription preinitiation complex. To assess the control these specific elements exercise on transcription bursting, we distinguished between two groups of genes based on the presence or absence of various core promoter elements (TATA box, Initiator element, CAAT box or GC box) within their promoter regions as described above and examined whether the two groups differed in transcription burst dynamics. In agreement with previous studies, we found that genes with promoters containing a TATA box had a significantly larger burst size than genes without (Figure 5C). The initiator element (INR), which also helps to recruit TFIID, was also significantly associated with a larger transcriptional burst size but here there was a comparably strong association with mean expression level (Figure 5C). Neither the CAAT box, nor the GC box were significantly associated with any of the three parameters (transcription burst size, burst frequency or expression level, Figure 5C).

### Epigenetic chromatin modifications associated with transcription burst kinetics

We next wanted to test the hypothesis that specific epigenetic marks correlate with transcription burst size or burst frequency. To achieve this, we analysed previously published ChIP-sequencing data for the MCF-7 cell line ^14,41,42^ and assigned ChIP-seq peak -log_10_(q-values) (FDR adjusted p-values) to corresponding genes. We then calculated the Spearman correlation of ChIP-seq -log_10_(q-values) and our transcription burst parameters (Figure 6A). All correlations were significant, again due to the large number of data points (genes) included in the statistical test. Generally speaking, correlations between the ChIP-seq data and transcription burst size or steady state expression level were stronger than for transcription burst frequency. The strongest correlations with burst size were seen for RNAPII (and its Ser5 and Ser2 phosphorylation), H3K27ac and H3K79me2. H3K27ac appeared to be the epigenetic mark most strongly correlated with transcription burst size (Spearman’s correlation coefficient, rho = 0.44), and was only slightly less correlated than Ser2 and Ser5 phosphorylation (rho = 0.42 and 0.43 respectively). Indeed plotting transcription burst size and burst frequency versus the H3K27ac ChIP-seq -log_10_(q-values) across all genes revealed the strength of this correlation (Figure 6B). Other epigenetic marks strongly correlating positively with transcription burst size include H3K9ac and H3K36me3, supporting their role as transcriptional activation marks (Supplementary Figure 8). The only epigenetic marks negatively correlating with transcription burst size were H3K27me3, H3K4me1 and H3K4me2 similarly supporting their usually repressive role in transcriptional regulation, however these correlations were not particularly strong. Other stronger correlations were seen between specific activating epigenetic marks, for example between H3K4me3 and H3K27ac, H3K9ac and H3K14ac (all at approximately rho = 0.4-0.7). KDM5B, which has recently been shown to drive variability in gene expression and phenotypic heterogeneity in ER α-positive breast cancer ^14^, showed no correlation with transcriptional burst frequency, but a positive correlation with burst size, indicating that this increase in KDM5B mediated phenotypic heterogeneity may be partially driven by increasing transcription burst size (Figure 6A, C).

**Figure 6.**
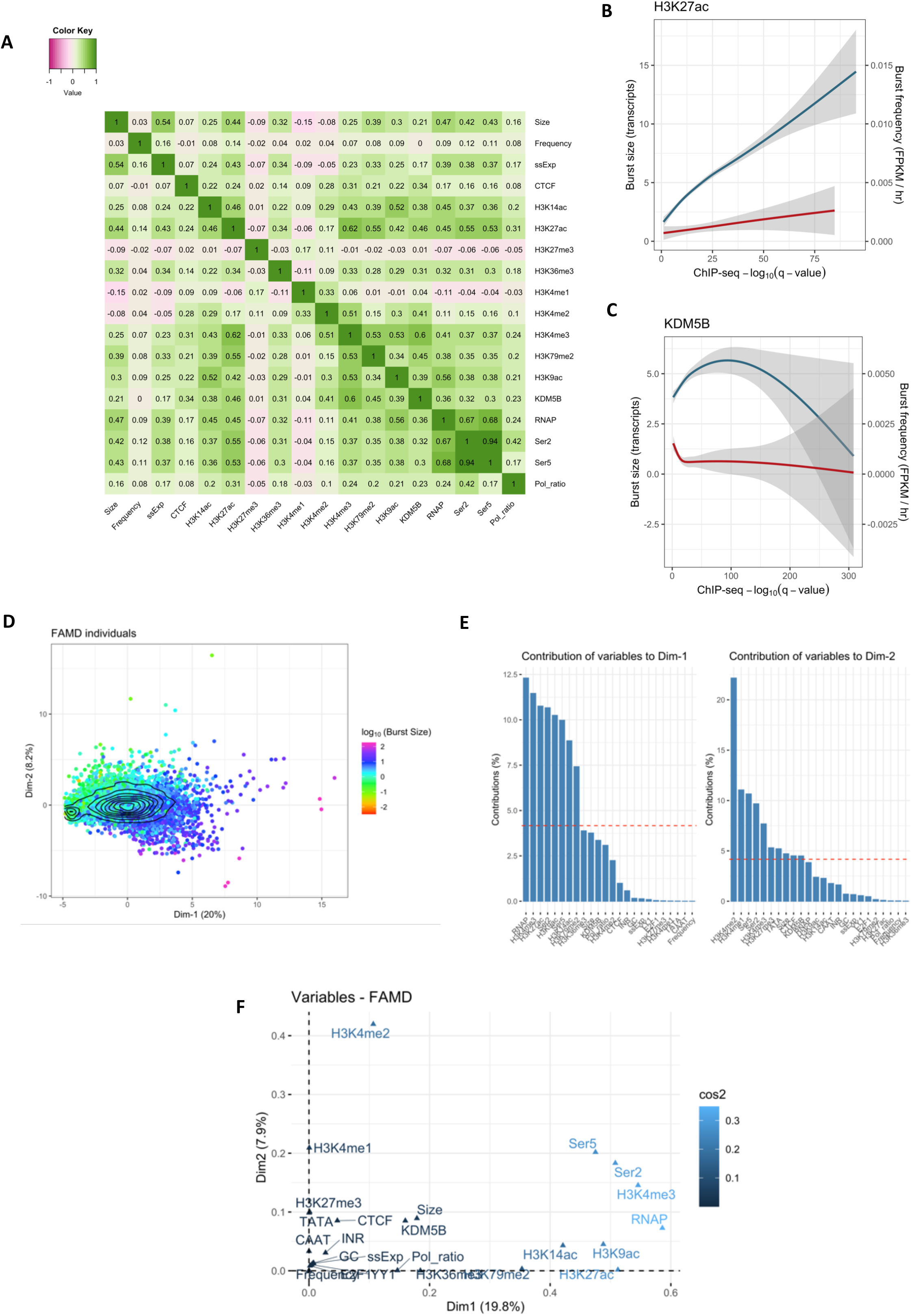
Many epigenetic features associate with transcription burst size while few associate with burst frequency. **(A)** Heatmap of Spearman’s correlation coefficients between transcription burst size, burst frequency, steady state gene expression (ssExp) level and all epigenetic and transcription regulatory factors analysed. Each individual correlation was significant owing to the large number of genes analysed (maximum number of 5,084 genes). **(B)** H3K79me2 ChIP-seq -log_10_(q-values) plotted against the generalised additive model (GAM) smoothed average for transcription burst size (blue) and burst frequency (red). Shaded regions show the 95% confidence interval. **(C)** As described above, for KDM5B transcription factor occupancy. **(D)** Individual genes plotted according to the first two dimensions following FAMD. Genes are coloured by burst size. **(E)** Contribution of individual variables to the first (left) and second (right) FAMD dimensions. Red dotted lines show the expected contribution for variables assuming equal contribution. **(F)** Squared loading plot depicting the association of each variable with the first two FAMD dimensions. Variables are coloured by the sum of their squared cosine (cos2) values of the first two dimensions, that is, the proportion of variance in a variable explained by the first two dimensions combined.

To identify any underlying relationship between transcription bursting parameters, the ChIP-seq -log_10_(q-values) and promoter motifs, we performed Factor Analysis of Mixed Data (FAMD ^43^), an extension of PCA which can handle both quantitative data (transcriptional parameters and ChIP-seq -log_10_(q-values)) and qualitative data (presence/absence of promoter motifs) simultaneously. Plotting the genes based on the first two dimensions of the resulting FAMD (which together account for almost 30% of the variance in the data) revealed two groups of genes (Figure 6D), corresponding to genes for which there are many assigned ChIP-seq peaks (right group) and genes for which there are few (left group). Transcription burst size correlated most strongly and positively with the first dimension, while it correlated more weakly and negatively with the second dimension. Analysis of the contribution of each variable to the first two dimensions revealed that the first dimension is dominated by RNA polymerase II binding and epigenetic marks known to be activating, whereas the second dimension is strongly (∼33%) comprised of H3K4me1/2 (Figure 6E). Plotting these variables in 2D space reveals that some marks, such as H3K79me2 and H3K27ac are strongly associated with the first dimension and independent of the second dimension, while H3K4me1 is exclusively associated with the second dimension (Figure 6F).

As observed above, many genes deviate significantly from the trend line between transcription burst size and mean mRNA expression level. We hypothesised that the relationships between specific epigenetic marks, transcription factor occupancy and transcriptional bursting parameters may be different for those genes for which the transcription burst size is considerably larger or smaller than expected based on the steady state gene expression. Analysis across the full dataset may obscure this information. To assess this aspect, we selected the genes with a larger burst size than expected from the mean expression level (n=173), genes with a smaller burst size than expected (n=282), and genes which lie along the trend line (n=441, Figure 3B). The correlation plots obtained after FAMD on these three separate groups indeed showed substantial differences between these groups. For the low burst size and trend line group, the burst size was predominately associated with gene expression level (ssExp) and RNAPII occupancy (RNAPII, Ser2, Ser5; Figure 7A), while epigenetic marks and the occupancy of KDM5B/CTCF appeared to play a minor role in explaining the variance in the data. Only for the large burst size group were the epigenetic marks associated with the transcription burst sizes (Figure 7A, right panel). To test the significance of these associations, Spearman’s correlation test was performed to quantify the extent to which each ChIP-seq dataset correlates with the burst size for the three groups of genes (Figure 7B, Supplementary Figure 9). Most epigenetic marks were more strongly correlated to transcription burst size at the higher end of the burst size spectrum rather than at low burst sizes (e.g. H3K9ac, H3K14ac, KDM5B). Transcription burst frequency was moderately negatively correlated to transcription burst size for the small burst size subset of genes only, suggesting that when the burst size is small given the expression level there is a coupling between the burst size and burst frequency. This coupling disappears in the other burst size groups. The same analysis was performed by dividing genes into three groups based on transcription burst frequency (Figure 3D, Supplementary Figure 10) but here correlations were much weaker.

**Figure 7.**
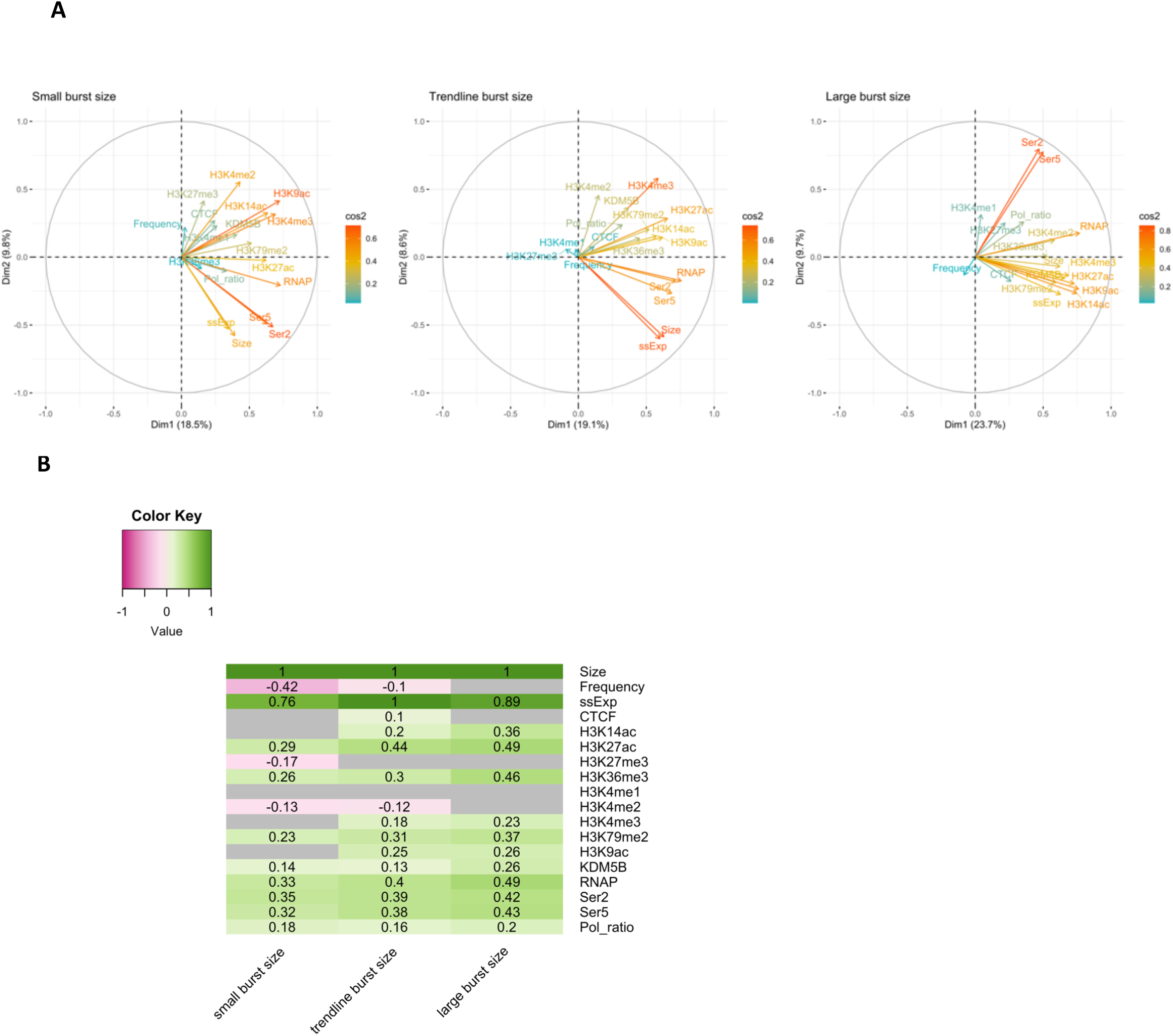
Large transcription burst sizes rather than small burst sizes are epigenetically defined. **(A)** Correlation circle plots for the small burst size (left, n = 282), trend line (middle, n = 441) and large burst size (right, n = 173) gene groups. Genes that exhibited a transcription burst size 5-fold less or 5-fold greater than expected based on their steady state gene expression were classified as small or large burst size genes, respectively. Genes with a burst size ± 0.1 fold of the expected burst size were categorised as trend line genes. Variables are coloured by the sum of their squared cosine (cos2) values of the first two FAMD dimensions, i.e., the proportion of variance in a variable explained by the first two dimensions combined. **(B)** Spearman’s correlation between transcription burst size and each epigenetic factor for each of the three gene groups described. Non-significant correlations are shaded grey. Values represent Spearman’s rho value from the correlation.

### Transcription burst size is associated with immediate responsiveness to tamoxifen treatment

It is frequently stated that transcription bursting allows for a rapid response of cells to changes in their environment, since few or short lived extra- and intracellular signals become amplified to produce many copies of mRNA. Our finding that genes with larger burst sizes are enriched for transcriptional regulators supports this idea, but we wanted to test this specifically and analyse the impact of applying environmental stress, such as breast cancer endocrine treatment. We analysed previously published triplicate RNA-seq datasets from MCF-7 cells treated with the selective estrogen receptor modulator (SERM) tamoxifen, with samples taken following 3, 6, 9, 12 and 24 hours of exposure ^44^. As above, two groups of genes were selected based on their transcription burst size (Figure 3B). The absolute fold change in gene expression of treated versus non-treated cells was calculated for each of the genes in the sets at each timepoint following tamoxifen treatment (Figure 8). On average, genes with large transcription burst sizes showed a 1.5 fold change at 3 hours of exposure, whereas those in the small burst size group only showed a 1.2 fold change. The gene expression fold change for the two gene sets subsequently converged somewhat until 9 exposure hours, after which the gradient of change became equal for the two groups. This data is in line with the notion that larger burst sizes facilitate a rapid response to environmental cues such as breast cancer endocrine treatment with tamoxifen.

**Figure 8.**
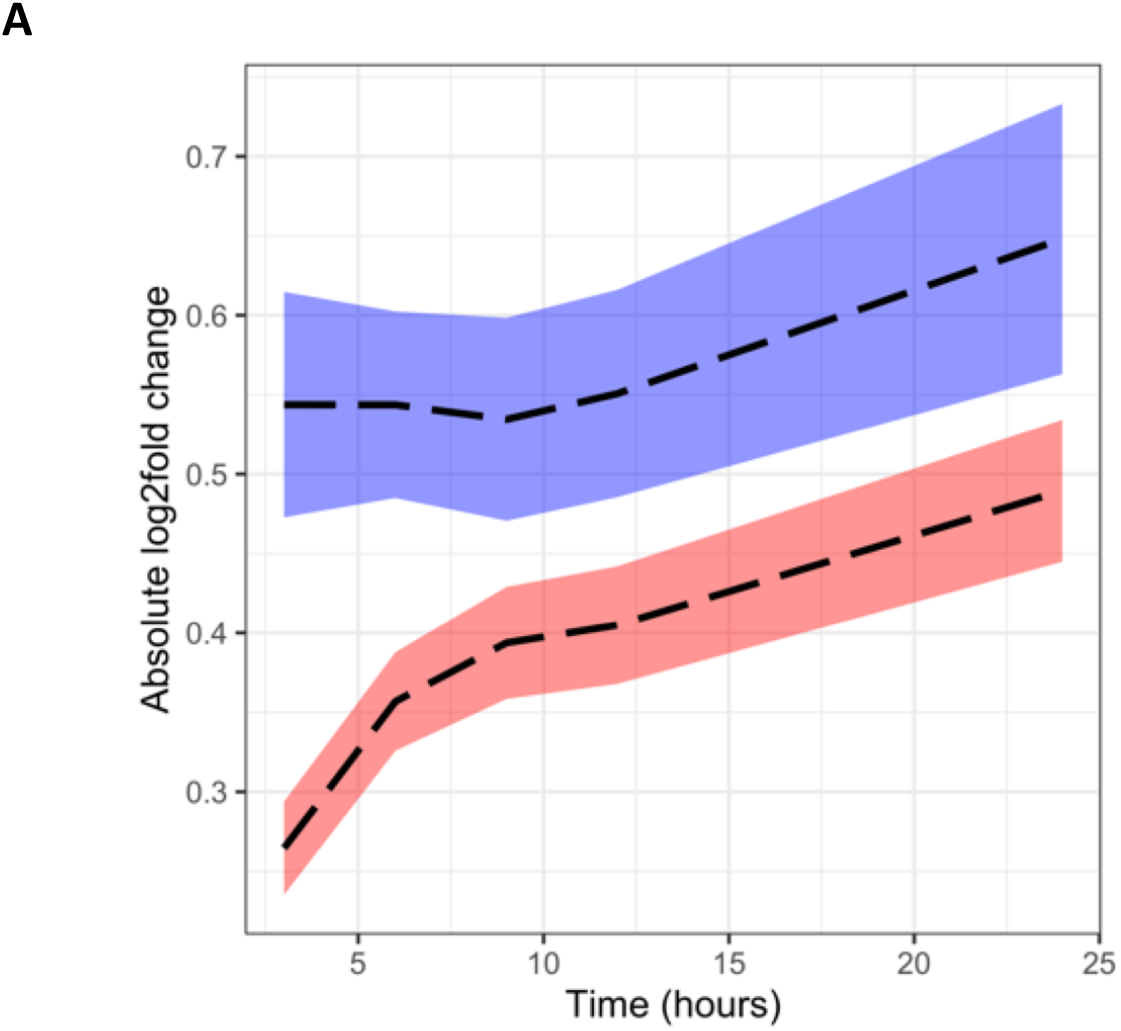
Large transcription burst sizes facilitate rapid gene expression changes in response to tamoxifen treatment. Triplicate RNA-seq data from MCF-7 cells treated with tamoxifen for 24 hours were downloaded and processed. Samples were taken after 0, 3, 6, 9, 12 and 24 hours of exposure. Dotted black lines represent mean absolute log2 fold change for the large burst size (blue, n = 173) and small burst size (red, n = 282) groups of genes, compared to the 0 hour sample (untreated). Differential gene expression was performed at the transcript level using featureCounts and DESeq2 packages. Shaded area shows the 95% confidence interval.

## Discussion

Our PP7 live cell imaging experiments have established a new way of quantifying burst sizes based on synchronisation of transcription across the cell population. While the main benefit of using such live cell systems lies in the ability to quantify transcription in single cells, synchronisation permits measurement of transcription burst sizes through bulk cell analysis, eliminating confounding factors such as static genetic heterogeneity and reliance on single cell sequencing and low throughput imaging. We calculated approximately 10 transcripts per burst for *GREB1* which is in accordance with previously published findings of between 5 and 17 transcripts ^29^, depending on the estrogen concentration within the media, and matched our average burst size of 12 transcripts calculated from peak detection of single cell PP7-*GREB1* transcriptional trajectories and the burst size of 10.2 calculated in the nascent RNA FISH experiment pre-length adjustment, or 5.8 post-length adjustment. Our live cell imaging data show that neither transcription burst size nor burst frequency was significantly affected by DRB treatment, validating the DRB removal strategy of our genome-wide transcription bursting measurements.

The *GREB1* transcription site intensity based on PP7-tagged transcript measurements in real-time was maximal at 50 minutes following DRB removal. This time window might integrate the time it takes for CDK9 to phosphorylate RNAPII CTD, for the thereby activated RNA polymerases to escape pause release and elongate to the site of PP7 loop integration, and for PP7 coat-GFP fusion proteins to bind to the emerging PP7 RNA loops. Given that the PP7 loops are integrated in the second exon of *GREB1* 17kb downstream of the TSS, we calculated that the RNA polymerases elongate along this region at approximately 0.34 kb/min. This is slower than previous estimates in between 1 and 6 kb/min^45–47^ and could reflect the fact that RNA polymerase elongation rates are slowest at promoter proximal regions, increasing along the gene body^48^. This slower elongation rate could also indicate that the accumulation time for PP7 coat-GFP fusion proteins is non-negligible or that there is residual DRB present.

Our 4sUDRB-seq nascent transcription data show that the vast majority of genes exhibit a characteristic burst in RNA production at transcriptional onset during a distinctive ON period, followed by a period of relative inactivity. This is in agreement with previous studies investigating genome-wide transcriptional dynamics which suggest transcription bursting is a ubiquitous feature ^1,49^. Indeed, only 687 genes failed to meet our criteria of transcription “burstiness” and even this could simply reflect a lack in sampling resolution or time course duration. The relationship between the transcription burst size and the steady state expression level of genes predicts heterogeneity in gene expression, revealing that genes involved in TNFA signalling via NFKB are particularly heterogeneous, while genes involved in fatty acid, glycolysis and cholesterol metabolism are more homogeneous. Unlike studies inferring transcription burst parameters from single cell transcriptomics, our approach of measuring transcription burst size from bulk RNA sequencing filters out any underlying genetic (static) differences between cells and so allows for the inference of more dynamic gene expression heterogeneity.

We define transcription burst size at the macro level, being the total number of transcripts produced between ‘deep repressive OFF states’, but being comprised of smaller microbursts, interspersed by ‘shallow OFF states’ (Figure 9). High frequency of microbursts results in a large macroburst size, and it is this macroburst that produces the large and intense transcription sites that we observed in RNA FISH data, driving non-Poisson distributed gene expression and heterogeneity. Our estimated transcription burst size for *TFF1*, of approximately 71 transcripts, was also in good agreement with previous studies using MS2 loop integration and live cell imaging, which suggested a macroburst size of 65 transcripts, itself being composed of smaller microbursts of approximately 2 transcripts ^32^. For a small gene of only 4.4 kb, it is possible to detect microbursts with MS2 or PP7 integration systems as only one or two RNA polymerases actively elongate along TFF1 at any given time, but transcription should continue for a longer period to produce a macroburst of approximately 60-70 transcripts, which is then often followed by a longer period of ‘deep repression’. For longer genes (such as *GREB1*), the residence time of nascent transcripts is much longer and the microburst resolution is lost. The quantified burst sizes here correlate with those measured by nascent RNA FISH. Differences between the nascent RNA FISH burst sizes and the true burst sizes could be due to these different residence times of nascent transcripts, mostly dictated by gene length, impacting the size and intensity of transcription sites in RNA FISH data but not affecting 4sUDRB-seq results. The multiscale model of transcription would also satisfy the prerequisites of the clock model of transcription, which we have previously shown to be able to produce such waves of mRNA production following synchronisation ^28^.

**Figure 9.**
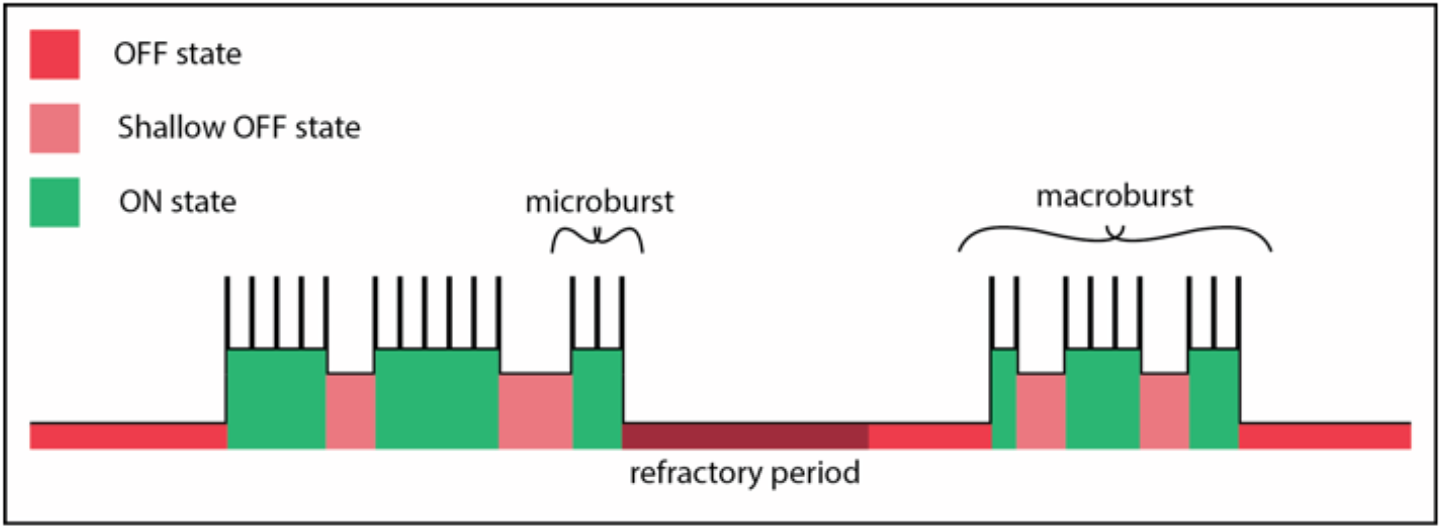
Schematic of the three-state model of transcription bursting in line with. ^32,64^. Promoters are capable of switching from deep OFF states into broadly defined ON states, which are themselves comprised of periods of active mRNA production (microbursts), and periods devoid of mRNA production (shallow OFF states). The sum of the mRNA produced during these broad ON states is the macroburst size which we have quantified in the present study using 4SUDRB-seq. The refractory period follows an ON to deep OFF state transition^27^. We propose that as a genome-wide trend the frequency of macrobursts contributes minimally to the steady state gene expression level, but that the macroburst size (i.e. the size times frequency of microbursts) is the key driver.

Intronic RNA is co-transcriptionally spliced and degraded ^34^. Our observation of intronic RNA decay within minutes following its production is in agreement with this. For larger genes this becomes less plausible since the size of the first intron could be exceedingly large (for example 300kb in the case of *RBFOX1*). However, recent evidence suggests that introns are actually removed in continuous chunks rather than in one large enzymatic step at exon flanks ^49^, which could explain why for these longer genes we still see rapid decay of intronic RNA.

YY1 binds to the promoters and enhancers of active genes and facilitates promoter-enhancer looping ^39^. Recent studies have suggested that enhancer-promoter contacts increase burst frequency but not burst size ^50^. This may depend on resolution and terminology, however, with respect to microburst and macroburst. We found no correlation between the presence of YY1 binding motifs in the promoter, and gene expression levels. However YY1 was the only transcription factor motif significantly associated with transcription burst size for both functional enrichment analysis methods employed, suggesting a role for YY1 in promoting larger transcription bursts, leading to higher levels of gene expression noise, at largely constant gene expression level. Given that YY1 is more strongly enriched at super enhancers relative to traditional enhancers ^51^ we propose a mechanism of transcription whereby YY1 mediated recruitment of distal super enhancers promotes large transcription burst sizes, possibly due to individual RNAPII recruitment from each of the enhancers within the super enhancer, or by preventing promoter state switching to the deep repressive OFF state. YY1 may also promote larger transcription bursts due to its ability to bind DNA and RNA simultaneously ^52^ and thereby cause a positive feedback on transcription initiation. Indeed blocking RNA production has been shown to reduce YY1 binding at promoters ^38^. Our result could also explain that in a study of phenotypic heterogeneity in 47 primary and metastatic estrogen-receptor (ERα)-positive breast cancer clinical specimens, the presence of YY1 was found to be a critical determinant of transcription output, mediating resistance to endocrine treatment ^15^.

The TATA box and Initiator element were significantly associated with larger transcription burst size, whereas no motifs were associated with transcription burst frequency. Presence of the TATA box has been repeatedly associated with larger transcription burst sizes and increased noise. Indeed mutational screening in yeast has proven the latter ^40^. We find that the association between the Initiator element and our transcription burst sizes was weaker than for the TATA box, and was comparable to the association between Initiator element and steady state gene expression level, suggesting that the Initiator element is less likely to be a driver of gene expression noise than of expression level. The presence of the CAAT box has also been suggested to drive burst size magnitude ^53^, but in the present study this effect was not significant (p-value = 0.051). However the CAAT enhancer binding protein beta (CEBPB) was one of the most significant transcription factors in our GAGE enrichment analysis. The GC box was not associated with any mean aspect of transcriptional kinetics.

Screening for epigenetic modifications correlating with our measurements of transcriptional burst size and burst frequency identified several associations. Histone acetylation is perhaps the best characterised epigenetic mark associated with regulation of transcription, specifically with transcription bursting ^29,54,55^. We find various histone H3 acetylation marks to be associated with elevated burst sizes such as H3K27ac and H3K9ac. Another epigenetic mark correlating with transcription burst size was H3K79me2, which was only slightly less correlated than RNAPII Ser2 and Ser5 phosphorylation. This appears to be in contrast to other findings in yeast, where knockdown of DOT1, the H3K79 methylation writer, produced higher levels of gene expression noise, ostensibly via a reduced transcription burst frequency ^56^. We found that correlations between epigenetic marks and transcription burst size were generally stronger when the burst sizes were larger than expected given the steady state expression level of the gene. This was not the case for burst frequency which was negatively correlated with burst size just for the small burst size group, which suggests differential coupling between the two parameters dependent on the magnitude of the burst size. The lysine demethylase KDM5B has previously been implicated in generating phenotypic and transcriptional heterogeneity in MCF-7 breast cancer cells ^14^. This is in agreement with our findings here, implicating enrichment at the promoters of genes with a larger burst size but not associating with increased burst frequency. It is therefore easy to envision how decreases in transcriptional variability arise as a result from inhibition of KDM5B, which improved the response of (ERα)-positive MCF-7 breast cancer cells to endocrine treatment ^14^.

Correlations between epigenetic marks and transcriptional burst frequency were not strong, only becoming more defined when looking specifically at the groups of genes with highest or lowest transcription burst frequency. At the low end of the transcription frequency spectrum, the factors most associated with frequency are H3K4me2/3 and H3K27ac, whereas at the higher end of the spectrum only H3K14ac is significantly correlated. From our attempt to identify the different components involved in transcription dynamics using FAMD, we noted that little of the variance in transcription frequency could be explained by the detected promoter motifs or enrichment of epigenetic components. Thus we conclude that control of the frequency of transcriptional bursting, at least at the macro level, is unlikely to be derived from these specific promoter elements or epigenetic histone modifications. Moreover, comparison of transcription burst frequency with the steady state gene expression level revealed a lack of association between the two (trend line gradient of 0.03), in contrast to burst size which is strongly correlated with steady state gene expression (trend line gradient of 0.84). A slight negative correlation between gene expression levels and RNA degradation rates accounts for this surplus of the expression level. Multiplying steady state expression level by the degradation rate, thereby adjusting for mRNA degradation rate differences between transcripts had a minimal effect on the trend line gradient (Δ=0.01), suggesting a minor role in dictating overall gene expression levels. Similarly, lack of change in the residual error of genes from the trend line implies that transcript degradation plays only a minor role in deviating genes from expected burst size-expression level coordinates. It is surprising that the transcription burst size contributes so much to the steady state gene expression level, but it may be that under varying conditions, when cells need to respond to stimuli, transcription burst frequency comes more into play.

Here, we have used the MCF-7 breast cancer cell line model to show that transcription bursting facilitates rapid gene expression changes in response to tamoxifen. However our methods and perhaps also our conclusions can be applied in a broader sense to interpreting the role of transcription bursting as a source of gene-expression noise, contributing to phenotypic heterogeneity and potentially driving pathological processes and drug resistance. We have uncovered correlative relationships between epigenetic factors and certain transcription factors with the magnitude of transcription burst size. We hope that our approach will allow for the investigation as to whether these correlative relationships between transcription bursting and the genomic/epigenetic composition are causal, for example by performing 4sUDRB-seq and quantifying macroburst sizes following epigenetic interference.

## Methods

### Transcription burst size measurements

We measured transcription burst size in three different ways: (i) Live cell imaging of PP7-tagged *GREB1* mRNA, (ii) 4sUDRB-seq, genome wide, and (iii) Single molecule nascent RNA FISH, for the genes *GREB1*, *AREG*, *PGK1*, *CD44*, *CD24*, *RAC1*, *EIF1* and *RACK1*. We will now detail these three methods.

#### PP7-tagged live cell imaging

The MCF7-GREB1-PP7 cell line was obtained from the Legewie lab ^29^ and grown in glutamax DMEM plus 10% foetal bovine serum and 1X Penicillin-Streptomycin-Glutamine. For imaging, cells were first washed with PBS, treated with 0.05% trypsin then plated onto an Ibidi polymer culture slide 24 hours prior to imaging. Cells were treated with 50 μM DRB dissolved in DMSO or DMSO only for 3 hours, then washed twice with 37°C PBS before the addition of 37°C imaging media (20mM HEPES pH 7.4, 137mM NaCl, 5.4mM KCl, 1.8mM CaCl_2_, 0.8mM MgCl_2_, 20mM glucose) supplemented with 1.25μL of 0.5mg/mL eGFP per mL of imaging media for burst size quantification. Cells were then selected and imaged every 5 minutes on a Leica SP8 confocal microscope at 512 × 512 × 22 resolution with a resulting voxel size of 0.36 × 0.36 × 0.4μm^3^.

#### 4sUDRB sequencing

MCF7-GREB1-PP7 cells were plated in 15cm culture dishes and grown to 80% confluency, then treated for 3 hours with 50 μM DRB (dissolved in DMSO) or an equal volume of DMSO. After 2 hours and 20 minutes, 100 μM of 4sU in DMSO was added to the control and 0 minute samples. For timecourse samples, media was then removed and cells were washed twice with 37°C PBS before fresh pre-warmed media was returned, containing 100 μM of 4sU in DMSO. Cells were then allowed to transcribe for 5, 10, 15, 20, 30 and 40 minutes. Following this, media from all samples was removed and 3mL of TRI reagent was added to lyse the cells and terminate transcription. RNA extraction was then completed following the TRI reagent protocol and final RNA pellets were dissolved in 100μL of RNase/DNase free water. For biotinylation, 100μL of 2.5x labelling buffer (25mM Tris HCL pH 7.4, 2.5mM EDTA) and 50μL of 1mg/ml EZ link HPDP-Biotin in DMF was added. Samples were then rotated at room temperature for 2 hours. To remove free biotin, 250 μL of chloroform:isoamylalcohol (24:1) was added and samples were vortexed for 30 seconds before being transferred to Maxtract tubes and centrifuged at 14,000g for 5 minutes at 4°C. The aqueous phase (∼200μL) was transferred to a fresh tube and 20μL of 5M NaCl, 200μL of isopropanol and 1μL of glycogen was added. Samples were vortexed and left at room temperature for 15 minutes then centrifuged at maximum speed (>20,000g) for 30 minutes at 4°C. The resulting RNA pellet was then washed twice in 80% ethanol before drying and dissolving in 100μL of RNase/DNase free water.

For pull down of biotinylated RNA, 50μL of Dynabeads MyOne Streptavidin T1 per sample was washed 3 times in 1X Binding and Washing buffer (5mM Tris-HCl pH 7.5, 0.5mM EDTA, 1M NaCl). Beads were then washed twice in Solution A (DEPC-treated 0.1M NaOH, DEPC-treated 0.05M NaCl) and once in Solution B (DEPC-treated 0.1M NaCl) then resuspended in twice their original volume (100μL per sample) of 2X Binding and Washing buffer. 100μL of beads was added to every sample before being rotated for 15 minutes at room temperature. RNA loaded beads were then separated on a magnet for 5 minutes and washed 3 times in washing buffer (100mM Tris-HCl pH 7.5, 10mM EDTA, 1M NaCl, 0.1% Tween 20) pre warmed to 65°C followed by washing once at room temperature. To release bound RNA, beads were incubated in 10mM EDTA, pH 8.2 with 95% formamide at 65°C for 10 minutes. Then samples were placed on a magnet and eluate was transferred to a fresh tube. Enriched RNA was then precipitated using isopropanol and 5M NaCl as described above, quantified using a Qubit RNA HS Assay Kit, and tested for quality on a Bioanalyzer with an RNA 6000 pico kit. The final yield of nascent RNA was approximately 0.2% of total input RNA taken for biotinylation.

200ng of nascent enriched RNA was sequenced using NEB ultra II directional RNA library prep kit in conjunction with NEBNext rRNA Depletion Kit. Samples were then checked for correct size on a Bioanalyzer. All 16 samples (two biological replicates) were sequenced on a mid-output v2.5 2×75bp (150 cycles) flow cell on the NextSeq 550 system.

#### Single molecule nascent RNA FISH

All exonic and intronic RNA FISH probe sets were designed using the Stellaris Probe Designer tool, with each probe set containing at least 48 individual unique probes. Exonic probes were conjugated with CAL Fluor® Red 590, whereas *GREB1* intronic probes were conjugated with Quasar® 670. 5 nmol of each probe set was resuspended in 400μL of TE buffer (10mM Tris-HCl, 1mM EDTA, pH 8.0) and stored at −20°C.

RNA FISH experiments were done following the standard protocol ^57^. Briefly, cells were transferred from culture to chambered polymer slides and allowed to adhere for at least 2 hours. Growth media was then gently removed and cells were washed with 1X PBS before being fixed for 10 minutes in 3.7% (vol./vol.) formaldehyde in 1X PBS. The cells were then washed twice with 1X PBS and permeabilised in 70% ethanol for at least 1 hour at room temperature. Following this, the ethanol was aspirated and 1mL of 1X Stellaris Wash Buffer A was added to the slide and incubated at room temperature for 5 minutes. Meanwhile, 100μL of hybridisation buffer was prepared (90μL Stellaris® RNA FISH Hybridisation Buffer, 10μL deionized formamide and 1μL of 12.5μM of probe solution). Wash Buffer A was then removed and the hybridisation buffer was evenly dispensed onto the slide. A glass coverslip was lowered onto the cells and the slide was placed into a small chamber alongside a tray of water to prevent the samples from drying. This chamber was then incubated in the dark for at least 4 hours at 37°C. The coverslip was removed and samples were incubated with 1X Stellaris Wash Buffer A for 30 minutes at 37°C, followed by another 30 minute wash at 37°C with 1X Stellaris Wash Buffer A containing 5ng/mL of DAPI. This was then aspirated and the samples were incubated with 1mL of 1X Stellaris Wash Buffer B for 5 minutes at room temperature, before being replaced by 2X SSC for imaging.

### Data analysis

#### Data analysis for PP7-tagged live cell imaging

Image analysis was performed using a custom semi-automated ImageJ script which allowed us to track and excise a 6×6 pixel region surrounding transcription spots. Voxels intensities of the resulting stacks were then summed and background was subtracted based on a region of the same size from the same nuclei. For molecule number quantification, an extracellular region was extracted and voxel intensities summed and divided by the expected number of GFP molecules within a given volume. Background subtracted transcription sites were divided by this intensity and then divided by 48 to reflect the fact that 48 molecules of GFP are able to bind one RNA molecule.

For single cell time-lapse analysis we applied a previously published peak detection algorithm ^58^ to the molecule numbers described above. We used the algorithm to calculate a moving mean and standard deviation based on every three timepoints, and set the threshold of detection to 3 standard deviations away from this mean. The frequency of bursts was simply the number of detected peaks per hour of time-lapse. Burst size was calculated by subtracting the initial number of molecules at the transcription site in every ON period from the maximal molecule number during that period. Burst sizes of zero were removed.

#### Data analysis for 4sUDRB-seq

Resulting FASTQ files were first aligned against the human genome (hg38) using BWA-MEM^59^, before being converted to SAM files and sorted using Samtools ^60^. To identify the transcription start sites for each gene, featureCounts from the Rsubread package was used at the transcript level and each gene was sorted based on transcript abundance. For each gene the transcription start site for the most highly expressed transcript was then taken and the others were disregarded. Visual inspection revealed that the *GREB1* transcription start site was incorrect and was therefore manually curated due to its importance in the study. A custom GTF annotation file was created for the 2kb intronic region downstream of every transcription start site with exons removed. Any genes with a size of less than 500bp after exon removal were discarded from the analysis. FeatureCounts was then run for every sample against this custom GTF file and the results were normalised by the total number of million mapped reads for each sample and by the size of the remaining intronic region in kb, to give FPKM. Biological replicates were then averaged and any genes showing 0 FPKM for any of the timepoints were removed (low expression filtering). The resulting time course data for every gene was smoothed using a loess smoothing function. Genes with maximal expression at 0 minutes were removed as were any genes that did not show an increase in expression at 5 minutes as compared to 0 minutes. To ascertain that the analysed genes had completed the transcriptional pulse in the 40 minutes time course, we removed any gene that did not show a decrease of at least 10% at any two sequential time points after 0 minutes.

Burst sizes were calculated for the remaining 8,429 genes by subtracting the expression at 0 minutes from the maximal expression at any point later in the timecourse. This burst size in FPKM was converted to absolute transcript number based on the quantified burst size for *GREB1* from live cell imaging of 10.33 transcripts and the 4sUDRB-seq *GREB1* burst size of 1.8 FPKM. The conversion factor between FPKM and number of transcripts for all genes is thereby taken to equal 5.7. Steady state gene expression level was determined by performing featureCounts on the full length transcripts for the control sample. Burst frequency was calculated as this steady state gene expression value in FPKM multiplied by the transcript degradation rate (from ^33^) and divided by the burst size in transcripts. The data units are therefore FPKM/h. Deming regression was performed for the steady state gene expression level and burst size or burst frequency, and groups of genes with large or small transcription burst sizes or burst frequencies were defined as those which deviated more than 5 fold from the expected value based on the regression. Defined gene groups were then tested for overlap with the MSigDB Gene Ontology collection using the online tool (www.gsea-msigdb.org/gsea/)

#### Data analysis for nascent RNA FISH

Samples were imaged on a Nikon Ti-E inverted confocal microscope with 60x oil objective in conjunction with Nikon NIS-Elements software. Samples were scanned simultaneously with 405nm and 561nm lasers for DAPI and CAL Fluor® Red 590 FISH probe respectively. For *GREB1* intron RNA FISH a 635nm laser was used. Optical sections in the z-direction were captured at 0.35μm intervals at a resolution of 512 × 512 pixels to give final voxel dimensions of 0.084 × 0.084 × 0.35 microns. 100 cells were imaged per gene per replicate. Image analysis was performed in ImageJ by manually selecting transcription sites based on maximal projections, followed by excision of the 12 x 12 pixel region (1.008μm^2^) across all z-slices. For each transcription site, the voxel intensities within this volume were summed. Single transcripts were selected from the nuclei and analysed in a similar way. The region excised around single transcripts was 5 x 5 pixels (0.175μm^2^) and the resulting voxel intensities were summed. Burst size for each transcription site was calculated by dividing the summed intensity of the transcription site by the average summed intensities of all single exon spots. We normalised transcription burst sizes by gene length by first averaging the gene lengths of all 8 genes, then dividing each gene length by this average length. Exon burst sizes were then divided by this normalisation factor to give length adjusted burst sizes.

### Functional enrichment and ChIP-seq analysis

Functional enrichment analysis was performed in R using the gage ^36^ and fgsea ^37^ packages. Genes were pre-ranked according to burst size and compared to MSigDB transcription factor or hallmark gene set databases. Gene sets containing YY1 binding factor and core promoter element motifs were downloaded from FindM ^61^. Significance between groups of genes based on these motifs was calculated using the Welch two sample t-test.

Raw FASTQ files for the ChIP-seq analysis were downloaded from GEO (accessions GSE104988, GSE23701 and GSE142011) using fastq-dump from the SRA toolkit. These raw files were then aligned to GRCh38 using BWA MEM ^59^ with default settings before being converted to BAM files using Samtools view ^60^. Peaks were called using MACS2 ^62^, with broad peak settings for histone modifications, and narrow peak settings for remaining data sets. To identify peaks that overlap with promoter regions, we created a bed file containing the regions between 1000 base pairs upstream and 100 base pairs downstream of every TSS in the analysis and identified overlap with ChIP-seq peaks using bedtools intersect command ^63^. For H3K36me3 we intersected with the full gene length since this is an epigenetic mark associated with the gene body. From the resulting bed file of peaks, we used the −log_10_(q-value) metric to perform FAMD and make figures. If any promoters/genes contained multiple peaks from the same ChIP-seq sample only the peak with the smallest q-value was taken. FAMD was performed in R using the FactoMineR and factoextra packages.

### Tamoxifen-treatment time-course data analysis

MCF-7 triplicate RNA-seq timecourse data was downloaded from ^44^. Samples were aligned to GRCh38 and converted to BAM files as described above, before performing featureCounts at the transcript level. Differential gene expression analysis was performed using DESeq2 with comparison of every timepoint to the control sample, and removal of any genes that were not significantly differentially expressed. Genes were grouped as described above i.e. based on their deviation from the expected transcription burst size or burst frequency relationship with the steady state gene expression.

## Data Availability

4sUDRB-seq data for all samples have been deposited on SRA under project number PRJNA691029. All other datasets used in this study are available from the authors upon request.

## Acknowledgements

We acknowledge Stefania Astrologo, Anchal Nigam, Misa Koncz, Diewertje Piebes and the EpiPredict consortium for their helpful suggestions. We thank Stefan Legewie (University of Stuttgart, DE) for kindly providing the MCF-7 GREB1-PP7 reporter cell line and Luca Magnani (Imperial College, UK) for exchanging experience and knowledge on breast cancer and endocrine resistance. We thank the sequencing unit and LCAM microscopy facility of the SILS for performing excellent services.

## Authors’ contributions

WFB performed the experimental and bioinformatic analyses, interpreted the results, generated figures, tables and drafted the manuscript. MALJ developed and performed image analysis of live cell PP7 data and nascent RNA FISH data. WFB and PJV conceived the study. PJV supervised the project and helped draft the manuscript. PJV, PDM and HVW interpreted the results and critically reviewed and co-edited the manuscript. All authors read and approved the final manuscript.

---

**Supplementary Figure 1.**
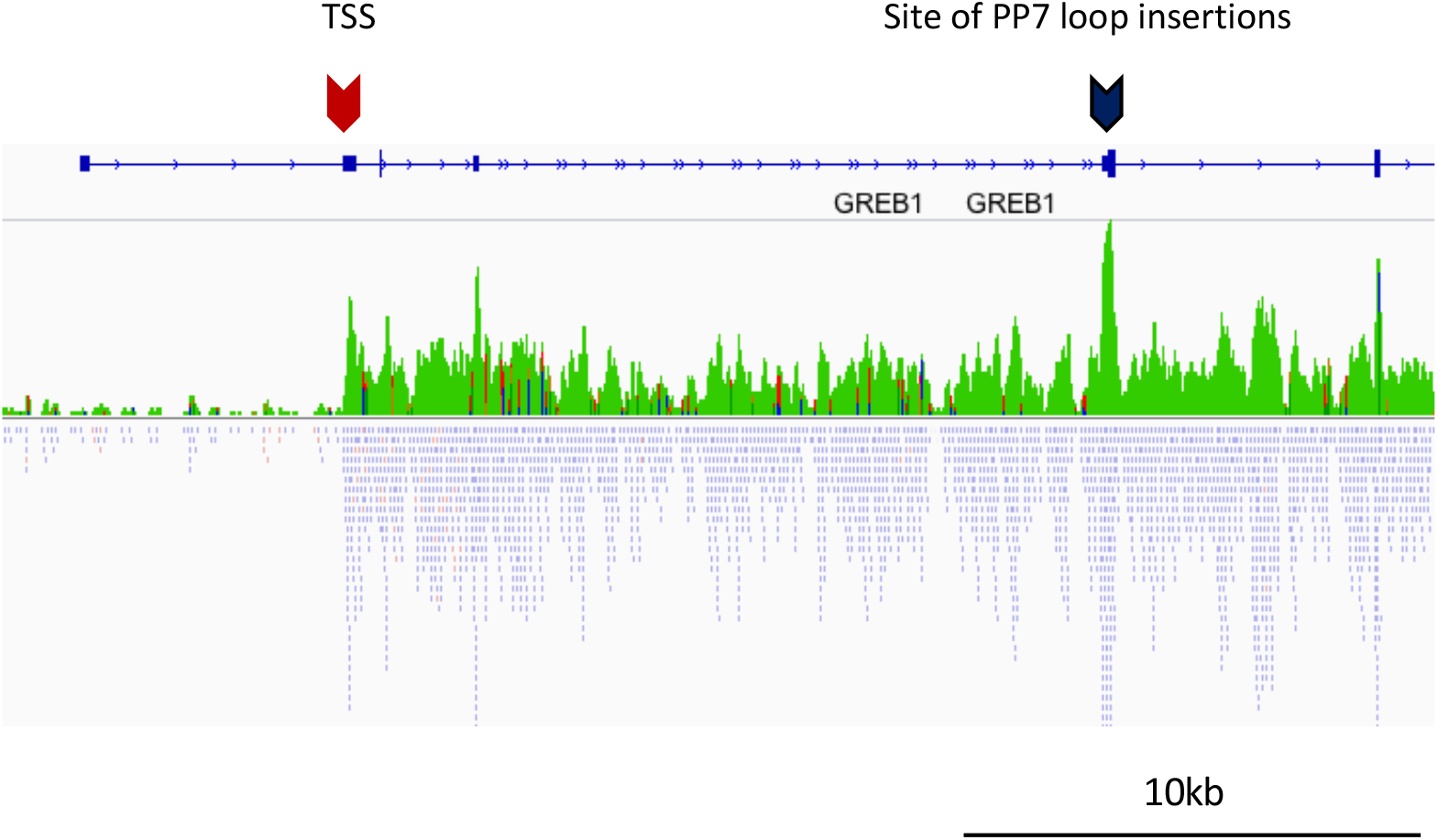
Illustration of the *GREB1* gene showing the site of PP7 loop integration in the second exon (black arrow). 24 PP7 stem loops were integrated, each of which may be bound by PP7-binding coat protein labelled with two GFP molecules (PCP□GFP). 4sUDRB-seq coverage (green) and reads (blue) for the control sample are shown below, highlighting the transcription start site (TSS) for *GREB1* (red arrow). Small blue arrows along the length of the gene represent the direction of transcription.

**Supplementary Figure 2.**
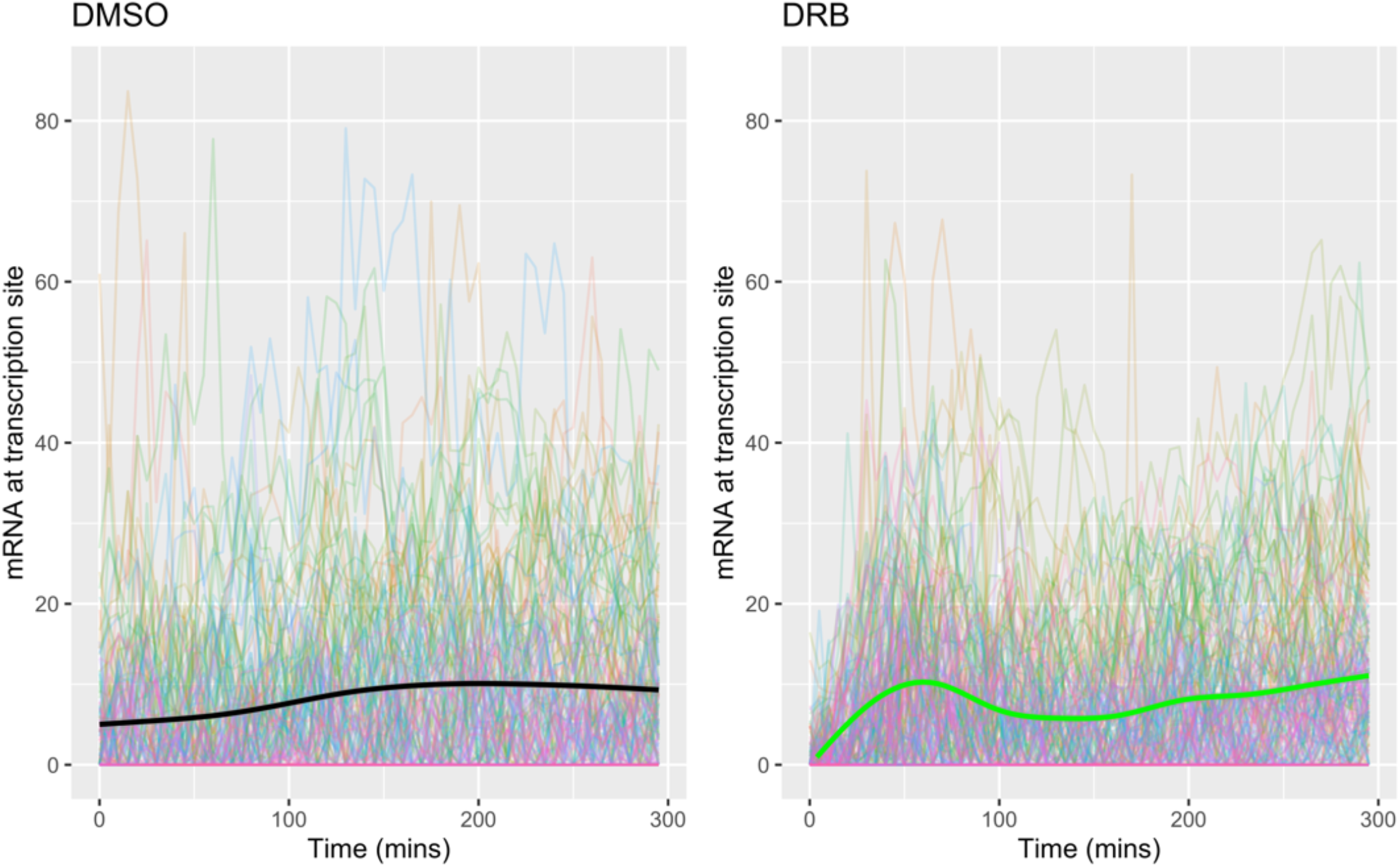
Transcriptional traces for all cells imaged following DRB treatment and release (right) or DMSO control (left). Thick black and green lines represent the average of the thinner individual traces of DMSO and DRB treated cells, respectively. Cells were imaged at 5 minute time intervals for 5 hours following DMSO treatment (left) or DRB treatment and release (right). The number of *GREB1* mRNA molecules at each transcription site was quantified by the addition of GFP at a known concentration to the imaging media (see Methods). The magnitude of the burst sizes between the two conditions is comparable.

**Supplementary Figure 3.**
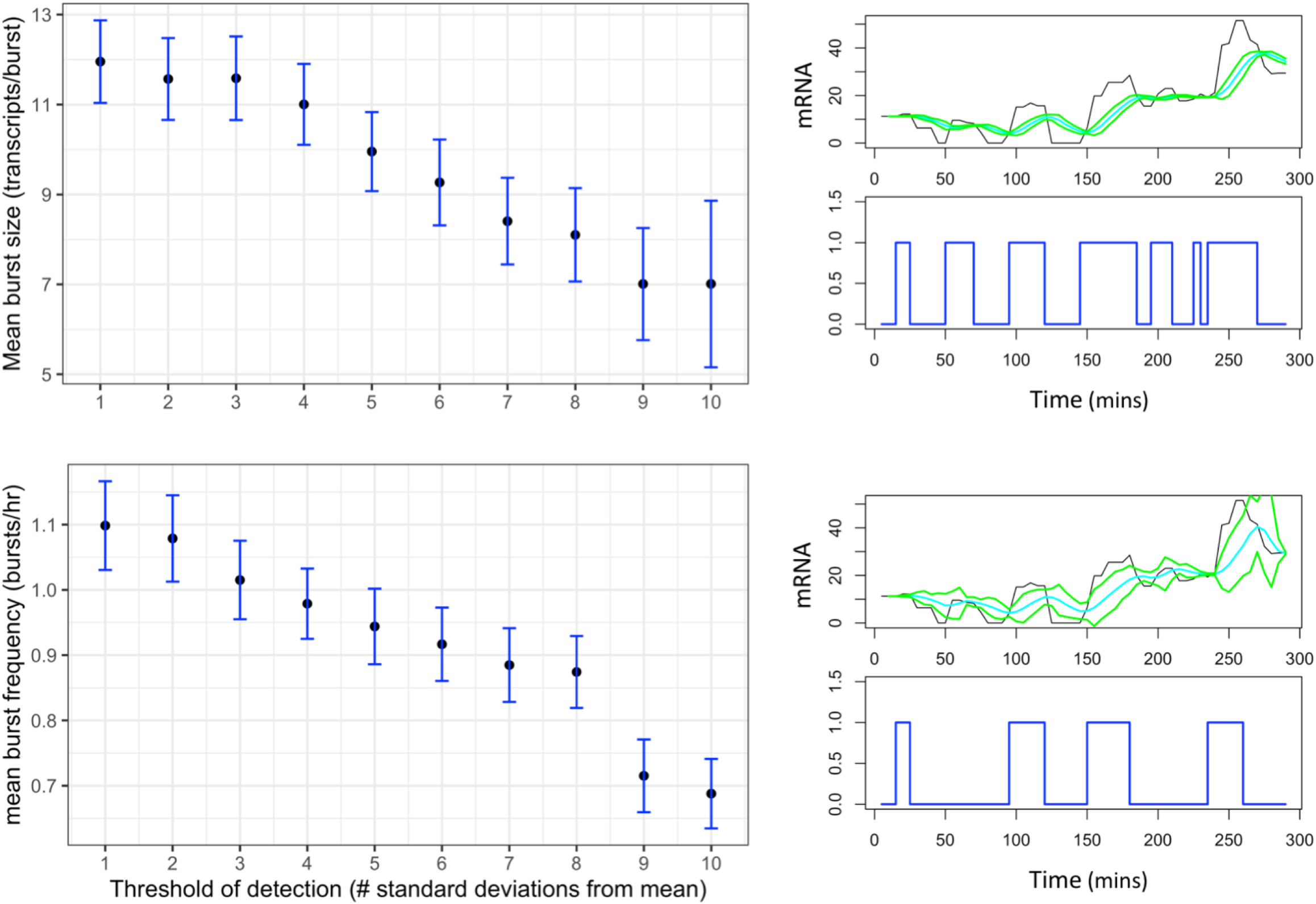
Increasing the threshold for peak detection reduces the mean transcription burst size (upper left) and mean burst frequency (lower left) from live cell *GREB1*-PP7 imaging trajectories. For these figures, all single cell trajectories were used, and the threshold of peak detected (based on the number of standard deviations from a rolling mean) was varied. Blue error bars show the 95% confidence intervals. An example of a single trace analysed using a threshold of 1 standard deviation from the mean appears to detect too many bursts (upper right). The same trace analysed with a threshold of 3 standard deviations is also shown (lower right). The raw time course data (black), is overlaid with a rolling average (blue) and standard deviations (green). Detected peaks are shown in the bottom panels (blue) as binary signals. For the analysis in Figure 1, a threshold of 3 standard deviations was chosen.

**Supplementary Figure 4.**
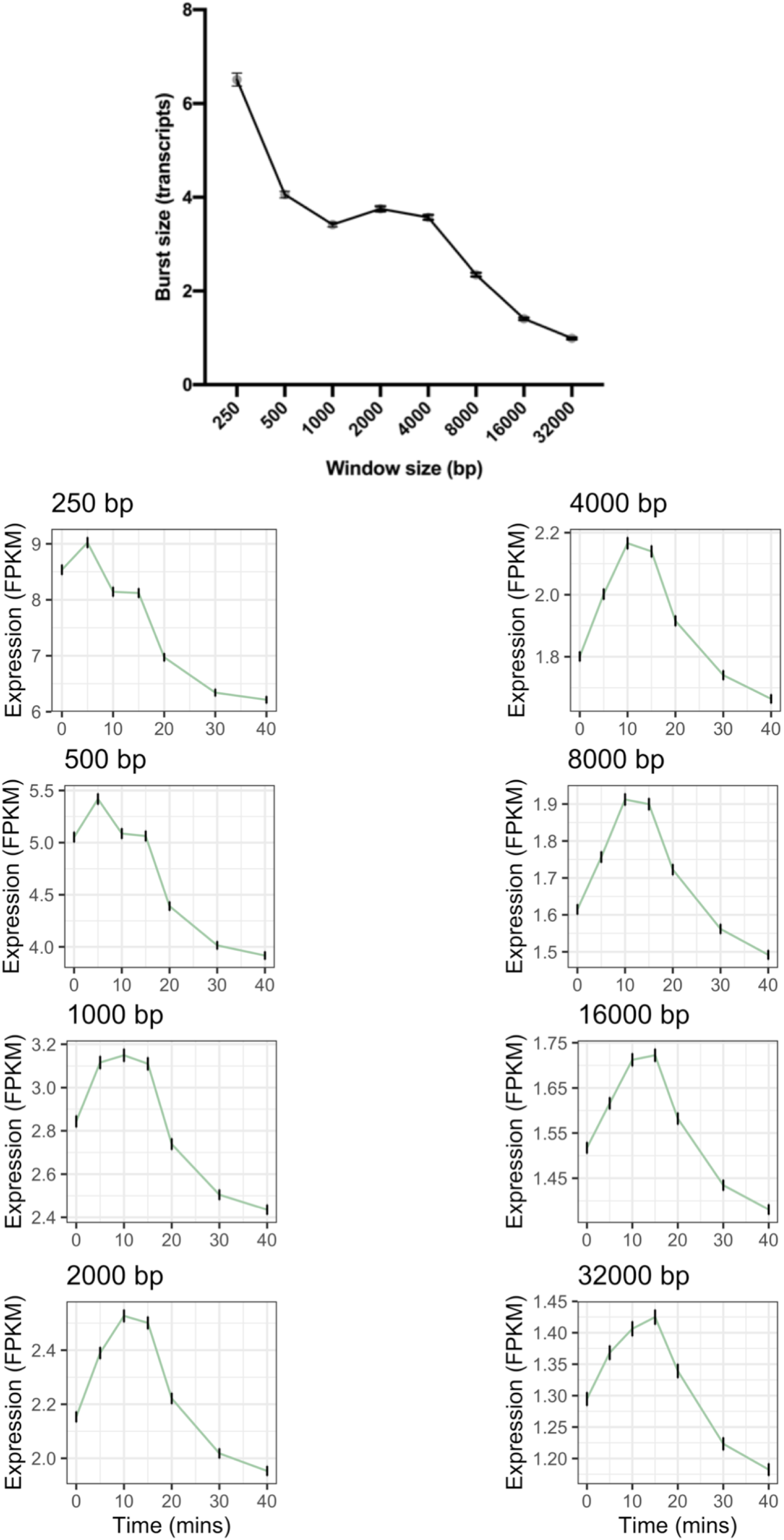
Genome-wide 4sUDRB-seq calculated mean transcription burst size generally decreases as the post-TSS window size for quantifying reads increases, with a plateau at 2000 bp. This window size was varied from 250 base pairs to 32,000 base pairs following gene transcription start sites, and the mapped reads within these regions were quantified and normalised by sequencing depth and remaining length after exon removal. The burst sizes were then calculated for each gene that passed low expression filtering (as described in Methods), and averaged (top). Error bars show ±SEM. Increasing this window size also delays the time of maximal transcription across the genome (bottom). Each plot shows the genome-wide average FPKM values mapped within the region after the transcription start site (250 bp, 500 bp etc). Error bars show ±SEM. Size filtering was not done here due to the differences in window size.

**Supplementary Figure 5.**
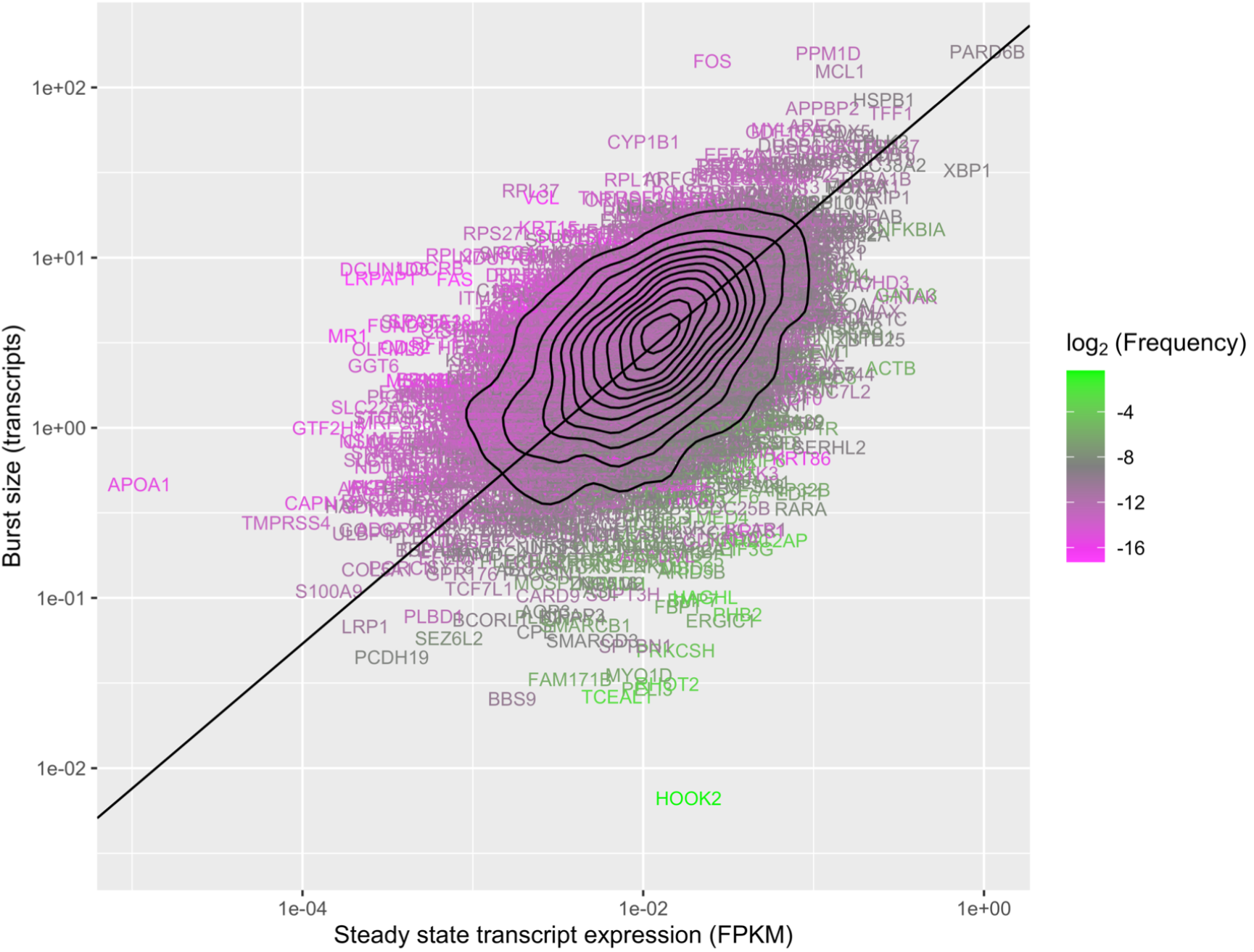
Adjusting for mRNA degradation rate differences between transcripts has a minimal effect on the trend line gradient and residual error. Deming regression (black solid line) of degradation adjusted-steady state gene expression level and transcription burst size for 5,084 genes. Genes are coloured by the log_2_(frequency). Adjusting for mRNA degradation rate differences increases the trend line gradient from 0.84 to 0.85, and decreases the residual error from 0.33 to 0.32. Degradation rates taken from ^33^.

**Supplementary Figure 6.**
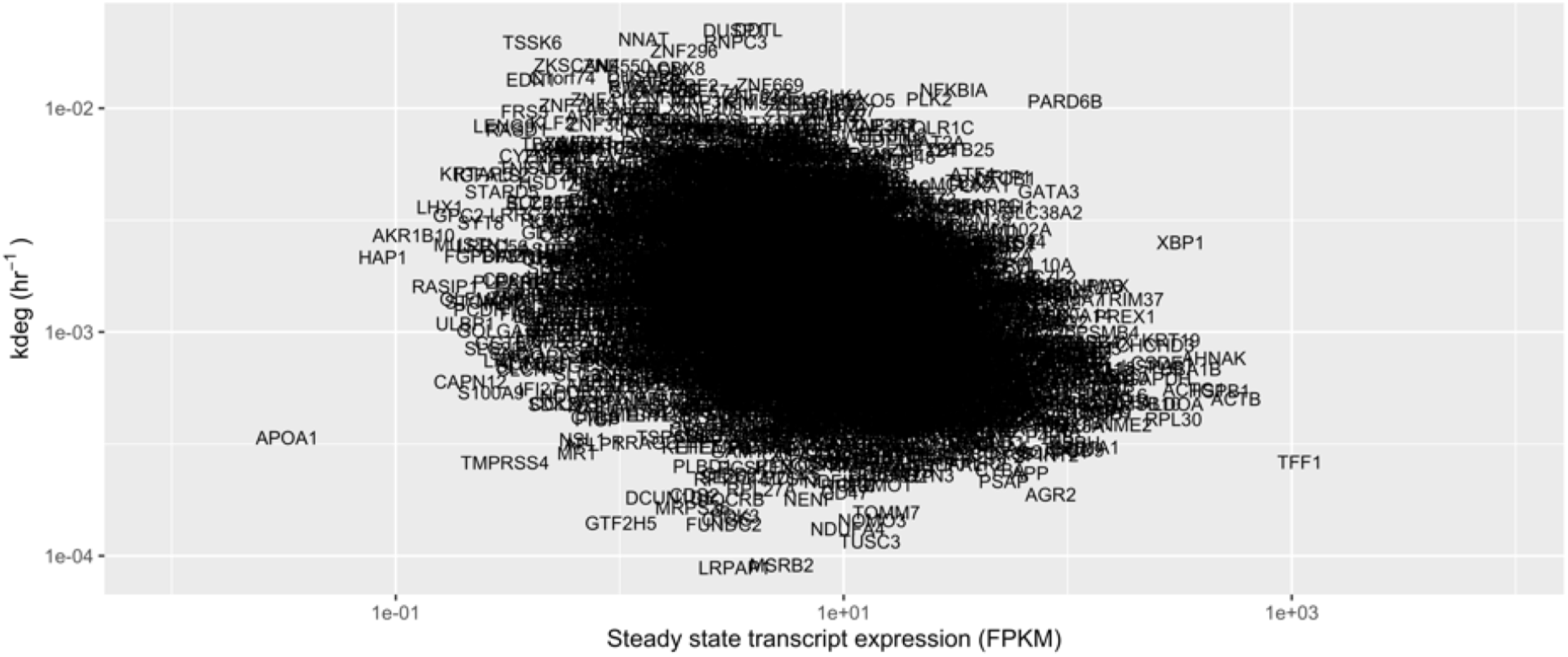
Genome-wide transcript degradation rates are negatively correlated to the steady state gene expression level. Transcript degradation rates (from ^33^) are plotted against the steady state gene expression level from 4sUDRB-seq control sample. The individual gene variance is large but there is a slight inverse correlation between the two parameters such that higher expression is associated with a reduced degradation rate.

**Supplementary Figure 7.**
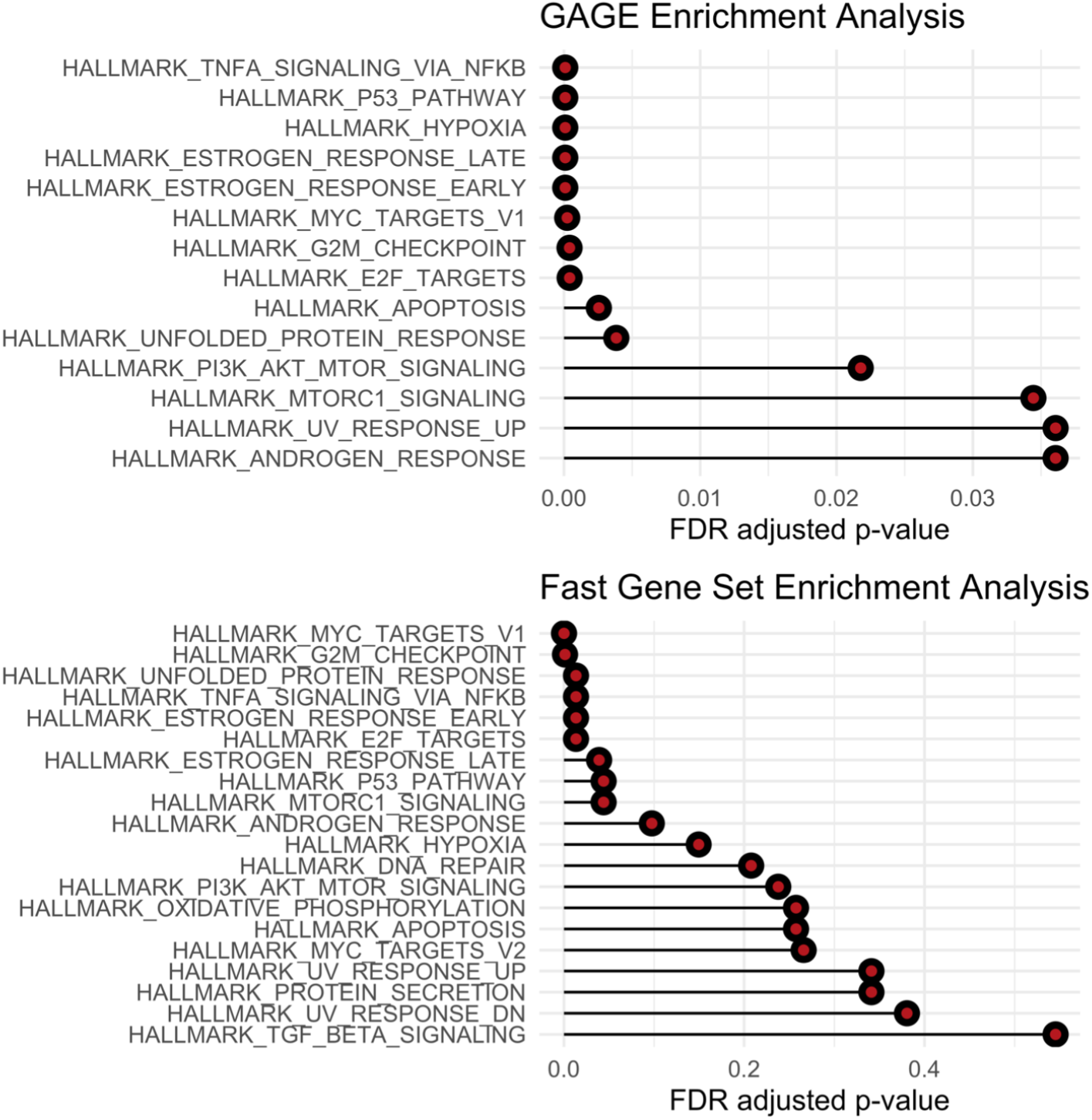
Pre-ranked functional enrichment analysis using the GAGE (top) and FGSEA (bottom) methods against the MSigDB Hallmark collection. 5,084 genes were ranked by their burst size and compared to the Hallmark collection. All displayed GAGE categories were significantly associated with a large burst size, whereas not all FGSEA categories are. The two subplots use different x-axes.

**Supplementary Figure 8.**
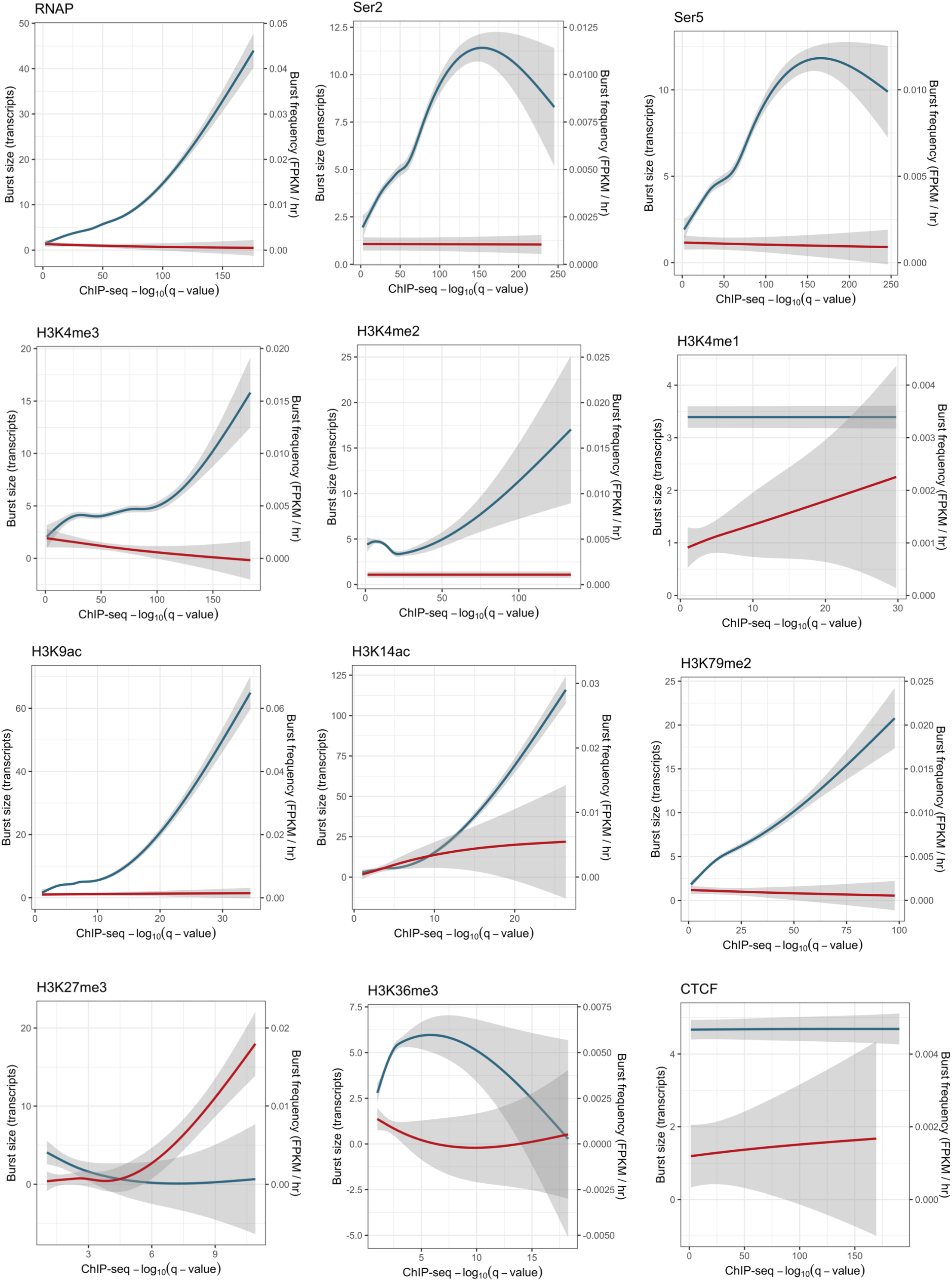
Correlations between ChIP-seq data and transcription burst size and burst frequency. Previously published ChIP-seq data was processed and peaks were assigned to overlapping gene promoters (except H3K36me3, see Methods). The matched ChIP-seq peak −log_10_(q-values) were then plotted against the generalised additive model (GAM) smoothed average for transcription burst size (blue) and burst frequency (red) for all epigenetic factors analysed. Shaded regions show the 95% confidence interval. The axes of each plot are unique.

**Supplementary Figure 9.**
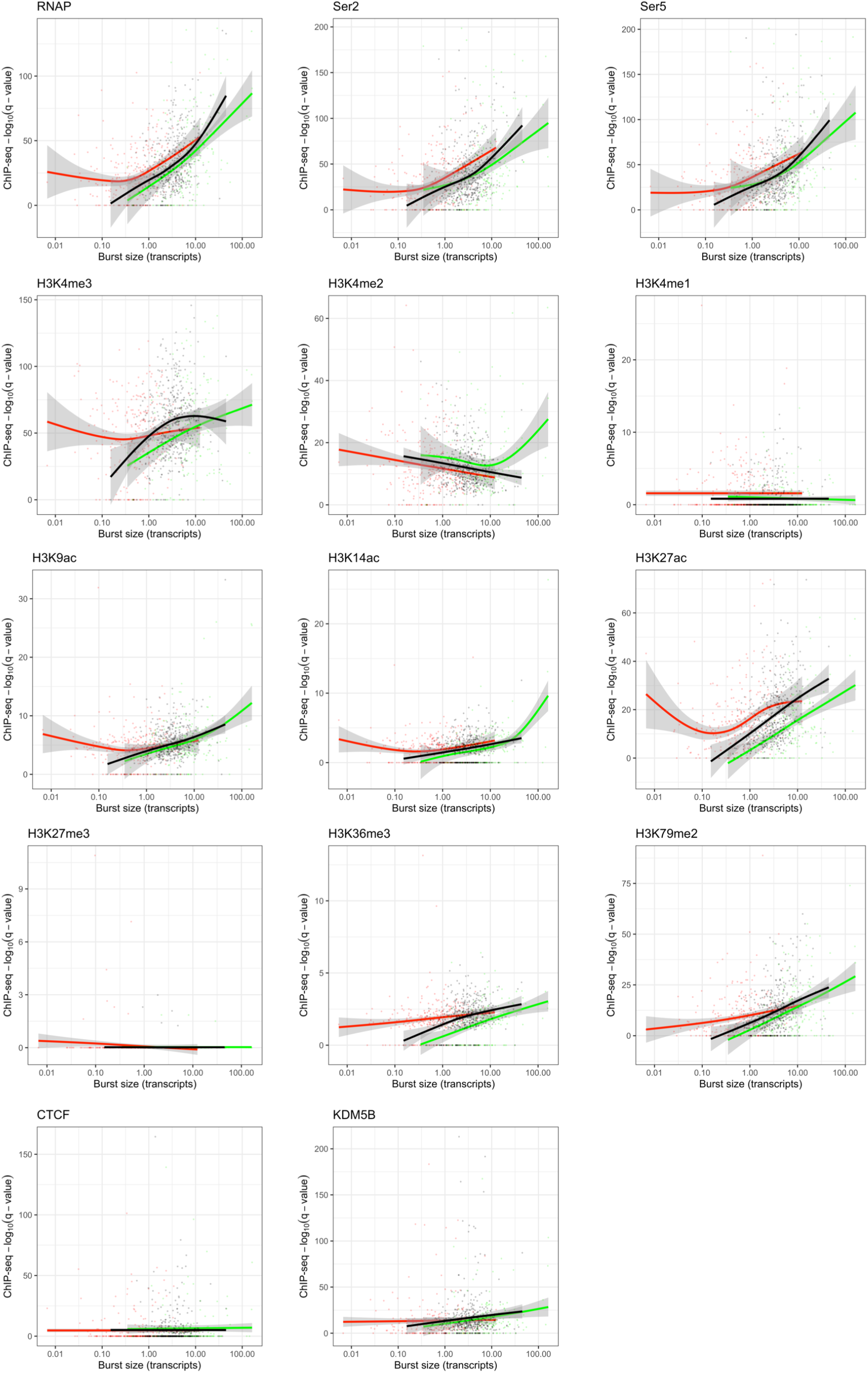
Correlations between ChIP-seq data and transcription burst size for each burst size group. Previously published ChIP-seq data was processed and peaks were assigned to overlapping gene promoters (except H3K36me3, see Methods). The matched ChIP-seq peak −log_10_(q-values) were then plotted against the generalised additive model (GAM) smoothed average. Genes belonging to the small burst size group, trend line group, and large burst size group are shown in red, black and green, respectively. Shaded regions show the 95% confidence interval. The axes of each plot are unique.

**Supplementary Figure 10.**
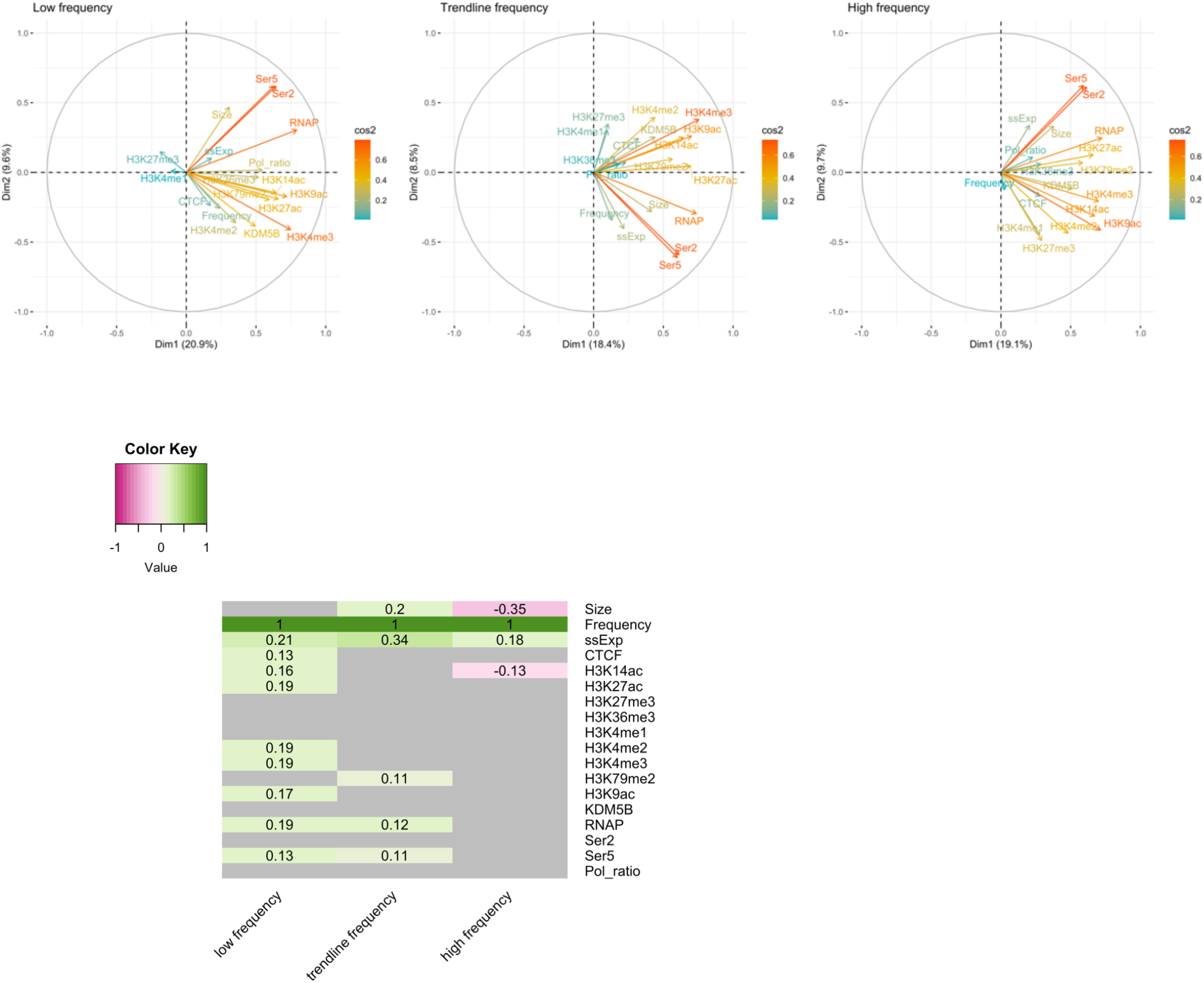
Correlation circle plots for the low burst frequency (left, n = 242), trend line (middle, n = 388) and high burst frequency (right, n = 348) gene groups. Genes that exhibited a burst frequency 5-fold less or 5-fold greater than expected based on their steady state gene expression were classified as low or high burst frequency genes, respectively. Genes with a burst frequency ± 0.1 fold of the expected burst frequency were categorised as trend line genes. Variables are coloured by the sum of their squared cosine (cos2) values, i.e., the proportion of variance in a variable explained by the first two dimensions combined (top). Spearman’s correlation between burst frequency and each epigenetic factor for each of the three gene groups described. Non-significant correlations are shaded grey. Values represent Spearman’s rho value from the correlation (bottom).

